# Dose-response functional and transcriptomic effects of follicle-stimulating hormone on *ex vivo* mouse folliculogenesis

**DOI:** 10.1101/2024.02.20.581188

**Authors:** Tingjie Zhan, Jiyang Zhang, Ying Zhang, Qingshi Zhao, Anat Chemerinski, Nataki C. Douglas, Qiang Zhang, Shuo Xiao

**Author notes:** Corresponding author: Shuo Xiao, Ph.D., 170 Frelinghuysen Rd, Rm 406, Piscataway, NJ, 08854, Qiang Zhang, M.D., Ph.D., 1518 Clifton Rd NE, Atlanta, GA 30322, USA. **Disclosure Statement:** The authors have nothing to disclose.

## Abstract

The gonadotropin-dependent phase of ovarian folliculogenesis primarily requires follicle-stimulating hormone (FSH) to support one or multiple antral follicles, dependent on the species, to mature fully, enabling ovarian steroidogenesis, oogenesis, and ovulation to sustain female reproductive cycles and fertility. FSH binds to its membrane receptor in granulosa cells to activate various signal transduction pathways and gene regulatory networks. Poor female reproductive outcomes can result from both FSH insufficiency owing to genetic or non-genetic factors and FSH excess as encountered with ovarian stimulation in assisted reproductive technology (ART), but the underlying molecular mechanisms remain elusive. Herein, we conducted single-follicle and single-oocyte RNA sequencing analysis along with other approaches in an *ex vivo* mouse folliculogenesis and oogenesis system to investigate the effects of different concentrations of FSH on key follicular events. Our study revealed that a minimum FSH threshold is required for follicle maturation into the high estradiol-secreting preovulatory stage, and the threshold is moderately variable among individual follicles. FSH at subthreshold, threshold, and suprathreshold levels induced distinct expression patterns of follicle maturation-related genes and the follicular transcriptomics. The RNA-seq analysis identified novel genes and signaling pathways that may critically regulate follicle maturation. Suprathreshold FSH resulted in multiple ovarian disorders including premature luteinization, high production of androgen and proinflammatory factors, and reduced expression of energy metabolism-related genes in oocytes. Together, this study improves our understanding of gonadotropin-dependent folliculogenesis and provides crucial insights into how high doses of FSH used in ART may impact follicular health, oocyte quality, pregnancy outcome, and systemic health.

## Introduction

Ovarian folliculogenesis is the process of development and maturation of ovarian follicles within the ovary. A dormant primordial follicle is activated and develops through the primary, secondary, antral, and preovulatory stages to reach full maturation. The early phase, from the primordial to early antral stage, is gonadotropin-independent (1–4). Despite gonadotrophin is not required for the follicular growth at this phase, FSH can still stimulate the secondary through early antral follicles to grow at a faster rate, therefore these more advanced follicles are gonadotrophin-responsive. In contrast, the late phase of folliculogenesis culminating in preovulatory follicles requires gonadotropins, especially follicle-stimulating hormone (FSH) produced by the anterior pituitary (5). In the late luteal and early follicular phase of each ovarian cycle, the regression of corpus luteum (CL) results in a remarkable decline of circulating inhibin, progesterone (P4), and estradiol (E2). As their inhibition on FSH is lifted, the rising FSH recruits a cohort of small antral follicles to grow. In many mammalian species, when the circulating FSH rises to a sufficient level, one or multiple antral follicles are selected to mature fully into preovulatory follicles while the remaining subordinate antral follicles regress (6–10). In monovulatory species, when FSH rises transiently above a threshold, a single antral follicle is selected as the dominant follicle. In polyovulatory species, multiple antral follicles, of similar nature to the dominant follicle in monovulatory species, are selected. The selected maturing follicles secrete hormones particularly E2 and support the oocytes within to acquire both meiotic and developmental competence (11,12). As the follicles continue to grow, the rising E2 stimulates the anterior pituitary to generate a surge of luteinizing hormone (LH) to trigger ovulation of a fertilizable oocyte or egg (1–4). The remaining follicular theca and granulosa cells luteinize to form a CL which primarily secretes P4 to prepare the endometrium for embryo implantation. These gonadotropin-dependent follicular events coordinately sustain female reproductive cycles and fertility.

At molecular levels, FSH binds to its membrane G-protein-coupled receptor, FSHR, in granulosa cells of early antral follicles to activate the canonical cAMP/PKA/CREB (cyclic adenosine monophosphate / protein kinase A / cAMP response element-binding protein) signaling pathway, which promotes the expression of a suite of genes crucial for terminal follicle maturation, such as cytochrome P450 family 19 subfamily A member 1 (*Cyp19a1*), LH/choriogonadotropin receptor (*Lhcgr*), and pregnancy-associated plasma protein A (*Pappa*) (13,14). *Cyp19a1* encodes the aromatase that converts theca cell-derived androgen to estrogen in granulosa cells (15). The induction of LHCGR in granulosa cells prepares preovulatory follicles to respond to the LH surge that triggers ovulation and luteinization (4,16). PAPPA acts as a secretory protease to breakdown insulin growth factor-binding proteins (IGFBPs) and free IGF which potentiates the expression of follicle maturation genes with FSH through the IGFR-PI3K/AKT/FOXO1 (phosphoinositide 3-kinases / protein kinase B / forkhead box protein O1) pathway (17–20).

Both insufficient and excessive FSH have been shown to cause poor female reproductive outcomes. FSH deficiency in humans owing to hypogonadotropic hypogonadism (HH) through either genetic (e.g., gonadotropin releasing hormone / GnRH or its receptor) or non-genetic (e.g., rapid weight gain or loss) reasons can result in delayed or absence of puberty, primary ovarian insufficiency (POI), amenorrhea, and/or infertility (21,22). In rodent models, deletion of FSH in gonadotrophs or FSHR in follicular granulosa cells leads to arrested folliculogenesis at the early antral stage, resulting in anovulation and infertility (1,2). Over the past few decades, recombinant human FSH (hFSH) has been utilized for controlled ovarian stimulation in assisted reproductive technology (ART) to simultaneously mature multiple follicles and collect multiple oocytes/eggs for *in vitro* fertilization (IVF) and embryo transfer, enabling successful conceptions in numerous otherwise infertile women (23). However, emerging epidemiological and experimental studies reported potential detrimental effects of high levels of FSH on the numbers of oocyte retrieved, oocyte quality, implantation/pregnancy success, maternal health, and health of the born offspring (24–31). In a heifer model, high doses of FSH induced diminished cumulus cell expansion, premature luteinization, and altered oocyte transcriptomic profiles (24). In cattle, the administration of high levels of FSH resulted in compromised E2 secretion and ovulation (25).

In women undergoing IVF, higher doses of FSH utilized to stimulate the reduced ovarian follicle pool characteristic of diminished ovarian reserve were associated with reduced numbers of retrieved oocytes and rates of clinical pregnancy and live birth (26–28); these clinical outcomes may reflect the diminished ovarian reserve or the higher doses of FSH administered (28,32). Women with “normal” ovarian reserve receiving high doses of FSH for infertility treatment have an increased risk of ovarian hyperstimulation syndrome (OHSS), a potentially life-threatening iatrogenic complication of ART characterized by ovarian swelling and leak of ovarian fluid into the peritoneal cavity due to increased capillary permeability and ovarian neo-angiogenesis (29,30). Together, these human and animal studies suggest that either insufficient or excessive FSH can be detrimental to ovarian folliculogenesis, oogenesis, and related reproductive outcomes. However, the molecular mechanisms underlying the differential responses and outcomes as stimulated by different FSH levels remain elusive.

We previously demonstrated that a 3D hydrogel encapsulated *in vitro* follicle growth (eIVFG) system faithfully recapitulates gonadotropin-dependent ovarian folliculogenesis, steroidogenesis, and oogenesis. Here, we used an *ex vivo* mouse folliculogenesis and oogenesis system along with single-follicle and single-oocyte RNA sequencing (RNA-seq) analysis and other approaches to investigate the effects of different concentrations of FSH on key follicular events. We hypothesize that FSH at subthreshold, threshold, and excessive suprathreshold levels induce distinct follicular transcriptomic responses and other follicular events, leading to different female reproductive outcomes. The novel findings of our study enable an in-depth understanding of gonadotropin-dependent ovarian follicle maturation, and the discovered molecular mechanisms provide crucial insights into how ovarian stimulation by high doses of FSH commonly used in ART may impact follicular health, oocyte quality, pregnancy success, and systemic health.

## Materials and Methods

### Animals

The CD-1 mouse breeding colony (Envigo, Indianapolis, IN) was maintained in the animal facility of Rutgers University under the controlled humidity and temperature environment within a 12-hour light-dark cycle. Mice were housed in polypropylene cages and provided with food and water ad libitum. Daily checks were conducted to accurately record the birth dates of female pups. Ethical approval for all animal studies was obtained from the Rutgers Institutional Animal Care and Use Committee (IACUC), in accordance with the NIH Guideline for the Care and Use of Laboratory Animals.

### Follicle isolation, encapsulation and eIVFG

Female mice on postnatal day (PND) 16 were euthanized using CO_2_, and ovaries were removed and collected in Leibovitz’s L-15 medium (Gibco, Grand Island, NY) supplemented with penicillin-streptomycin and fetal bovine serum (FBS, Sigma-Aldrich, St. Louis, MO). Ovaries were cut into 6-8 pieces and enzymatically digested in L-15 media containing Liberase TM (Roche, Indianapolis, IN) and DNase I (Worthington Biochemicals, Freehold, NJ) for 25 min. Multilayered secondary follicles with diameters ranging from 130-160 μm, and normal morphology were selected. 0.5% alginate hydrogel in PBS and calcium solution were then used to encapsulate a single follicle in one alginate bead. Encapsulated follicles were maintained in the αMEM Glutamax media (Gibco) with 1% FBS for 30 min. Follicles were then cultured in 96-well plates individually with 100 μL growth media (GM). The GM consisted of 50% αMEM Glutamax (Thermo Fisher Scientific, Waltham, MA) and 50% F-12 Glutamax (Thermo Fisher Scientific) supplemented with 3 mg/ml bovine serum albumin (BSA, Sigma-Aldrich), 1 mg/ml bovine fetuin (Sigma-Aldrich), 5 μg/ml insulin-transferrin-selenium (ITS, Sigma-Aldric), and varying concentrations of recombinant rFSH (gifted by Organon, Jersey City, NJ, USA). Encapsulated follicles were cultured under GM with FSH at 5 mIU/mL on day 0-4, then with FSH at 5, 10, 20 or 30 mIU/mL on day 4-6. Follicle images were captured using an Olympus inverted microscope every two days (Olympus Optical Co Ltd, Tokyo, Japan). Follicle size was determined by calculating the average of two perpendicular diameters using Image J software (National Institutes of Health, Bethesda, MD).

### *Ex vivo* ovulation induction

Following a 6-day *ex vivo* culture under GM with different concentrations of FSH, grown antral follicles were free from alginate encapsulation by digestion in an L-15-based lysis solution containing 1% FBS and 10 IU/mL alginate lyase (Sigma-Aldrich) for 20 minutes at 37°C. Follicles were then cultured in αMEM-based maturation media supplemented with 10% FBS and 1.5 IU/mL Human chorionic gonadotropin (hCG, Sigma-Aldrich) in a an incubator (37°C, 5% CO_2_, Thermo Fisher Scientific) for 14 hours to induce *ex vivo* ovulation. After hCG treatment, follicles were imaged to assess the rupture rate. “Ruptured follicle” was shown with broken follicular wall with an expanded cumulus-oocyte complex (COC), while “unruptured follicle” maintained an intact follicular wall. Oocytes were collected from ruptured follicles and imaged to evaluate oocyte meiosis by calculating metaphase II (MII) oocytes, identified by the presence of a polar body. Germinal vesicle (GV) or germinal vesicle breakdown (GVBD) oocytes were characterized by the absence of a polar body.

### Measuring hormones production in conditioned media

To examine the impact of different FSH concentrations on ovarian steroidogenesis, conditioned follicle culture medium was collected on day 4 and 6 before ovulation induction and day 9 after *ex vivo* ovulation. The concentrations of E2, P4 and testosterone (T) were measured using competitive enzyme-linked immunosorbent assay kits (ELISA, Cayman, MI, USA), based on the competition between the hormones and a hormone-acetylcholinesterase (AChE) conjugate. The dilution factors of conditioned culture medium for E2, P4, and T on day 4 and 6 were 50-fold, 400-fold, and 5-fold, respectively, and 400-fold for postovulatory P4 on day 9, to ensure the detection values within the standard curve range.

### Single follicle RNA extraction and reverse transcription-quantitative polymerase chain reaction (RT-qPCR)

Total RNA of a single follicle was extracted using Arcturus PicoPure RNA isolation kit (Applied Biosystems, Carlsbad, CA) following manufacturer’s instructions. cDNA was synthesized using the Superscript III First-Strand Synthesis SuperMix kit (Invitrogen, Carlsbad, CA) and stored at -80°C to prevent degradation. qPCR was conducted in a 384-well plate, with reaction mixtures comprising Power SYBR Green PCR Master Mix (Applied Biosystems), forward primer (0.25 μM), reverse primer (0.25 μM), and cDNA template. The qPCR was performed using the ABI ViiA 7 real-time PCR system (Applied Biosystems) with the following thermocycling program: 95°C for 10 min, 40 cycles at 95°C for 15 s and at 60°C for 1 min, followed by a melting stage at 63°C for 25 s. Glyceraldehyde-3-phosphate dehydrogenase (*Gapdh*) was used as an endogenous control for normalizing mRNA expression values of each gene, calculated using the 2^(-ΔΔCT)^ method. Primer sequences are provided in the Suppl. Table 1 (33). The qPCR assay was performed in at least 6 biological replicates with 2-3 technical replicates.

### Single-follicle RNA-seq and data analysis

Follicles on day 4 and 6 treated with FSH of 5, 10, 20 and 30 mIU/mL were collected for single-follicle RNA-seq analysis. Total RNA of a single follicle was extracted as described above. The cDNA library construction and low-input mRNA-Seq were performed on an Illumina NovaSeq PE150 platform by Novogene (Novogene Corporation, Sacramento, CA). High-quality paired sequencing reads were obtained with the effective rate of 96-99% and the error rate of 0.03%. The sequencing data were processed in the Partek Flow software for further analysis. Bowtie 2 module was employed to filter out the rDNA and mtDNA contaminants. The filtered reads were aligned to the whole genome of Mus musculus-mm10 assembly via HISAT 2. The aligned reads were quantified based on the Ensembl Transcripts release 99 annotation model using the Partek EM algorithm. Gene counts were normalized using the log2-transformed Transcripts Per Million (TPM) plus 1.Gene symbols were mapped to the HUGO Gene Nomenclature Committee (HGNC) to filter out pseudogenes. Differential gene expression analysis was conducted via the DESeq2, with differentially expressed genes (DEGs) identified with a fold change of either > 2 or < 0.5 and False Discovery Rate (FDR)-value < 0.05. Principal component analysis (PCA) was performed using the R package. Gene Ontology (GO) and Kyoto Encyclopedia of Genes and Genomes (KEGG) enrichment analysis were carried out using the Database for Annotation, Visualization, and Integrated Discovery (DAVID) (34,35).

### Single-oocyte RNA-seq and data analysis

MII oocytes were collected at 14 hours post-hCG for single oocyte SMART-Seq analysis. cDNA libraries were prepared using the Nextera XT Library Prep Kit. In brief, cDNA was synthesized, and amplified with the following thermocycling program: 95°C for 1 min, 10 cycles at 98°C for 10 s, 65°C for 30 s and 68°C for 3 min, and then 72°C for 10 min. cDNA was then purified with AMPure XP beads, and quality was examined using a DNA High Sensitivity Bioanalyzer. cDNA tagmentation was performed with Nextera XT Index Kit v2. cDNA library was further amplified with the following thermocycling program: 72°C for 3 min, 95°C for 30 s, 13 cycles at 95°C for 10 s, 55°C for 30 s and 72°C for 30 s, and then 72°C for 5 min. The sequencing of premade library was further performed on an Illumina NovaSeq PE150 platform by Novogene (Novogene Corporation, Sacramento, CA). Raw sequencing data were analyzed using the Partek Flow software as described above.

### Fuzzy clustering of transcriptomics data

To characterize the temporal and dose-dependent expression trajectories of genes, their expression values were analyzed using fuzzy clustering with the R package of M-fuzz (36,37). Expression levels of DEGs in follicles treated with FSH on day 4 and in follicles treated with FSH at 5, 10, 20, and 30 mIU/mL were combined. The M-fuzz workflow was applied to cluster gene expression trajectories across culture days from day 4 to 6 and FSH concentrations from 5 to 30 mIU/mL, generating 6 clusters. Genes (membership > 0.6) exhibiting different trajectories were further selected as input for GO enrichment analysis using DAVID.

### Statistical analysis

All data were shown as mean ± standard deviation (SD). Data were analyzed by one-way analysis of variance (ANOVA) followed by Tukey’s multiple comparison using GraphPad Prism (SPSS Inc., Chicago, IL) to analyze the significance of different FSH concentrations on follicle growth, hormone secretion, follicle rupture rate, and the expression of related genes. A two-sided *P*-value less than 0.05 was considered statistically significant and denoted with asterisks as *\*P<0.05; ** P<0.01; *** P<0.001*.

## Results

### A FSH threshold is required for gonadotropin-dependent follicle maturation

Immature mouse follicles at multilayered secondary stage were first cultured in eIVFG with 5 mIU/mL FSH from day 0 to 4 to mimic the gonadotrophin-responsive growth. Follicles were then switched to treatment with different concentrations of FSH, at 5, 7, 8, or 10 mIU/mL, from day 4 to 6 to determine the minimum levels of FSH required for terminal follicle maturation toward the preovulatory stage and related follicular functions. These include follicle growth and E2 secretion, two hallmark events of FSH-dependent follicle maturation (38). Treatment with FSH from day 4 to 6 stimulated follicle growth in a concentration-dependent manner (Figure 1A-1B). At 10 mIU/mL FSH, the terminal follicles on day 6 reached 352.68 ± 19.50 µm in diameter, which were significantly larger than the diameter of 319.57 ± 19.08 µm when follicles were cultured with 5 mIU/mL FSH (Figure 1A-1B). The diameters of follicles cultured with 7 or 8 mIU/mL FSH on day 6 fell in between, at 338.28 ± 15.23 µm and 339.99 ± 20.75 µm, respectively. When the increment of follicle diameter from day 4 to 6 was used as a metric for evaluating follicle growth, the increments were FSH concentration-dependent, and at 8 or 10 mIU/mL, FSH had significantly greater stimulatory effects than at 5 mIU/mL (Figure 1C). Notably, there were considerable inter-follicle variations within and between FSH concentration groups: some follicle diameter increments in the 7 or 8 mIU/mL FSH groups were on par with the 10 mIU/mL group.

**Figure 1.**
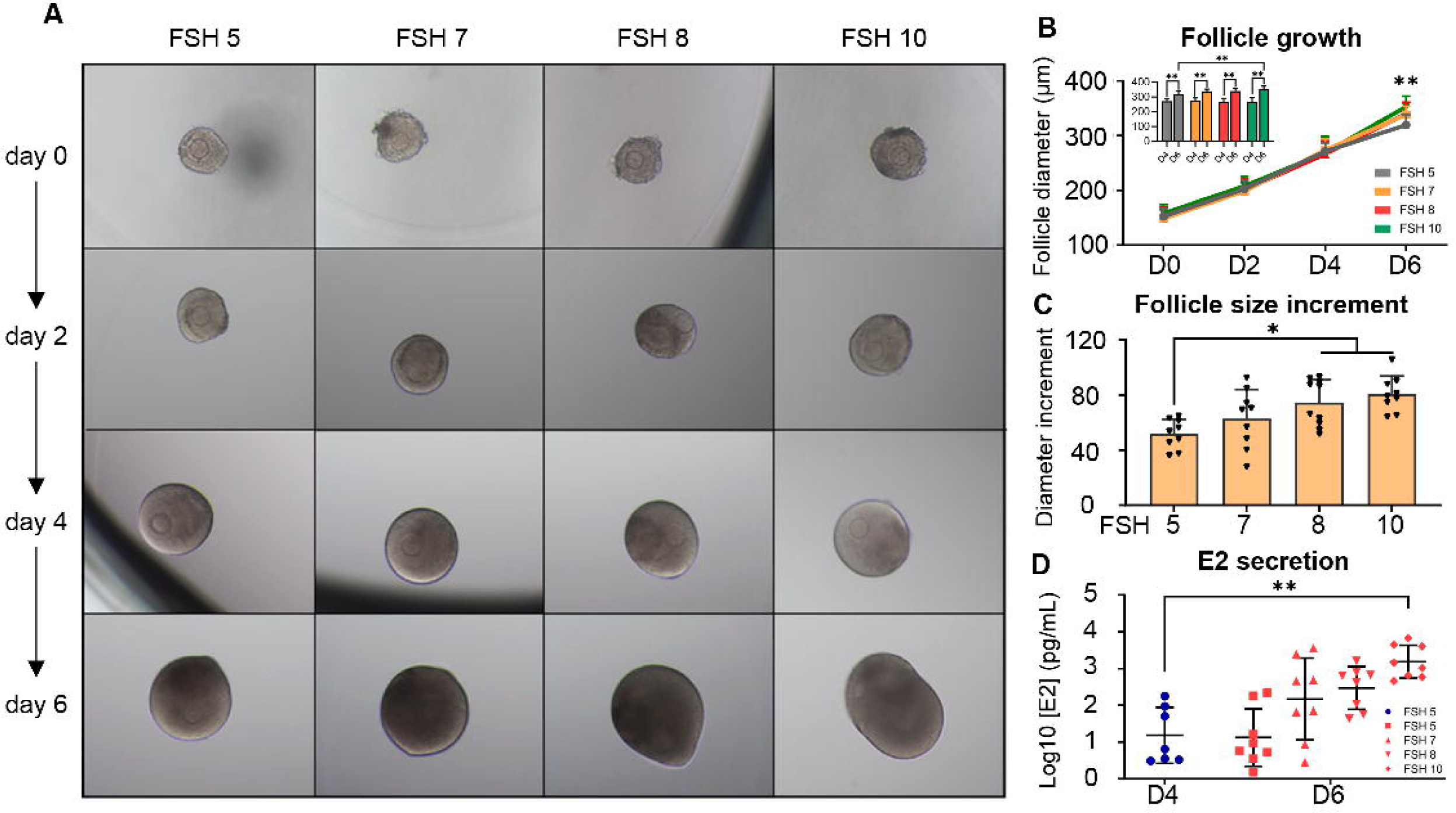
The effects of various FSH concentrations of 5, 7, 8, or 10 mIU/mL on mouse follicle development and hormone secretion in an *ex vivo* folliculogenesis system. Follicles were cultured for 4 days with FSH of 5 mIU/mL to grow to the early antral stage, and further treated with FSH at 5, 7, 8, or 10 mIU/mL from day 4 to 6. (A) Representative bright-field images of follicles on day 6. (B) Diameters of follicles cultured from day 0 to 6. (C) Change in follicle diameter, reported as the increment, from day 4 to 6 with treatment of different concentrations of FSH. (D) Concentration of estradiol (E2) in the conditioned culture medium on day 4 and 6. N = 8-15 follicles in each treatment group and 3 replicates were performed. Error bar: standard deviation; *\*P<0.05; ** P<0.01*.

There were also remarkable inter-follicle variations in the amount of E2 secreted, often across over one order of magnitude even within the same FSH concentration group (Figure 1D). Despite the significantly larger follicle size on day 6 than day 4 (Figure 1B insert), follicles cultured with 5 mIU/mL FSH secreted E2 on day 6 in a range that was comparable to day 4 (Figure 1D). At 7 or 8 mIU/mL FSH, some follicles switched to a much higher E2-secreting state on day 6, but statistically the average E2 secretions were insignificant compared to day 4 due to the heterogenous responses within groups (Figure 1D). At 10 mIU/mL, nearly all follicles secreted E2 at high levels on day 6, which were non-overlapping with day 4, and the average E2 secretion level was significantly greater, for about 100-fold, than day 4. Taken together, these results suggest that for mouse follicles cultured in eIVFG, a minimal FSH threshold, likely somewhere above 5 mIU/mL in the eIVFG system, is required to stimulate gonadotropin-dependent follicle maturation; moreover, this threshold is variable among individual follicles, likely ranging between 5 mIU/mL as the lower bound and 10 mIU/mL as the upper bound, and when FSH is at or above 10 mIU/mL, nearly all follicles would switch to a high E2-secreting state.

### Excessive suprathreshold FSH augments preovulatory secretion of P4 and T

Given that the FSH thresholds range between 5-10 mIU/mL for different follicles, we next examined whether excessive suprathreshold FSH concentrations alter key follicular events. Similar to the results in Figure 1, at 5, 10, 20, and 30 mIU/mL, FSH concentration-dependently increased the follicle terminal size on day 6 (Figure 2A-2B) and diameter increments from day 4 to 6 (Figure 2C). Although some follicles were lagging in growth in the 10-30 mIU/mL FSH groups, the average follicle diameters or increments in all 3 groups were significantly greater than the 5 mIU/mL group. Like in Figure 1D, 5 mIU/mL FSH did not cause any increase in E2 secretion between day 4 and 6, but 10 mIU/mL FSH significantly increased E2 secretion by about 11-fold with a nearly clean separation between the two groups (Figure 2D). Compared with 10 mIU/mL, 20 and 30 mIU/mL FSH further increased E2 secretion by about 3 and 2-fold, respectively, but the average increments were statistically insignificant with the majority of the individual follicle responses overlapping (Figure 2D). The amounts of P4 secreted by follicles treated with 5 mIU/mL FSH were comparably low on day 4 and 6 (Figure 2E). In contrast to E2 secretion where the effects of 5 and 10 mIU/mL FSH are nearly dichotomized, 10 mIU/mL FSH caused only negligible, statistically insignificant increase in P4 secretion compared with 5 mIU/mL (Figure 2E). However, 20 or 30 mIU/mL FSH significantly increased P4 secretion on day 6 by about 3 and 2-fold respectively compared with 10 mIU/mL (Figure 2E). The amounts of T secreted by follicles treated with 5 mIU/mL FSH were similar on day 4 and 6 (Figure 2F). FSH at 10 mIU/mL increased T secretion by 6-fold compared to 5 mIU/mL on day 6 but the increase was insignificant (Figure 2F). At 20 or 30 mIU/mL, FSH further stimulated T secretion to significantly higher levels than follicles on day 4 and follicles treated with 5 or 10 mIU/mL FSH on day 6 (Figure 2F).

**Figure 2.**
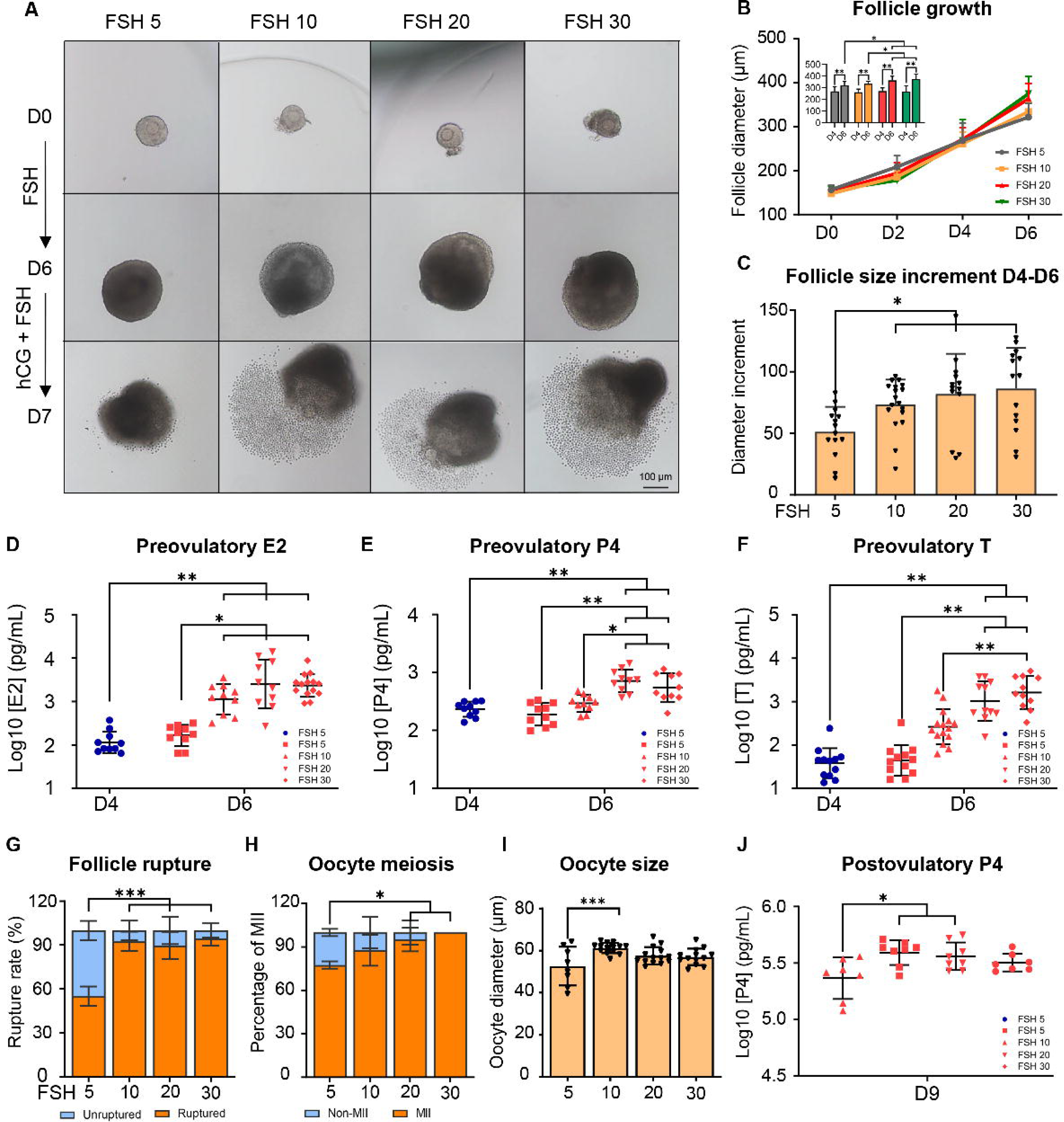
The concentration-response effects of FSH on mouse follicle growth, hormone secretion, ovulation, and luteinization in an *ex vivo* folliculogenesis system. Follicles were cultured for 4 days with FSH of 5 mIU/mL to grow to the early antral stage, and then treated with FSH at 5, 10, 20 or 30 mIU/mL from day 4 to 6. (A) Representative bright-field follicle images on day 0, 6, and 7. (B) Diameters of follicles culture with FSH on day 0-6. (C) The increment of follicle diameters on day 4-6 with treatment of different concentrations of FSH. Concentrations of E2 (D), P4 (E) and T (F) in the conditioned culture media on day 6. After the *ex vivo* hCG ovulation induction, follicles were further cultured for 48 hours for luteinization and P4 secretion. Percentages of ruptured follicles (G) and MII oocytes (H) with treatment of various FSH concentrations. (I) Diameters of oocytes from ovulated follicles. (J) Concentrations of P4 in the conditioned culture media on day 9. N = 8-15 follicles in each group and 2-3 replicates were included. Error bar: standard deviation; *\*P<0.05; ** P<0.01*.

We next examined various key ovulatory and post-ovulatory events with follicles on day 6 treated with hCG to induce ovulation *ex vivo*. For follicles on 5 mIU/mL FSH, 55.04 ± 6.59% of them were able to rupture, and among the ruptured follicles, 77.50 ± 2.50% of ovulated oocytes were at the MII stage, which were significantly lower than the 92.50 ± 6.61% follicle rupture rate and 87.78 ± 10.72% MII percentage in the 10 mIU/mL FSH group (Figure 2G-2H). Oocytes ovulated from follicles treated with 5 mIU/mL FSH were significantly smaller than the oocytes from follicles treated with 10 mIU/mL FSH (52.66 ± 9.18 µm vs. 61.28 ± 2.48 µm, Figure 2I). Compared to 10 mIU/mL, 20 or 30 mIU/mL FSH did not further increase the follicle rupture rates, MII oocyte percentages, and oocyte size (Figure 2G-2J). We then cultured post-ovulatory follicles for additional 48 hours to allow for luteinization and P4 secretion by the formed CL. ELISA results showed that compared to 5 mIU/mL, 10 mIU/mL FSH significantly increased P4 secretion, but 20 or 30 mIU/mL FSH did not further promote P4 secretion (Figure 2J). In summary, these results further confirmed that a minimal FSH threshold above 5 mIU/mL is required to stimulate terminal follicle maturation, as manifested by endpoints such as follicle growth and ovarian hormone secretion. For ovulation-related categorical endpoints such as follicle rupture, oocyte meiotic resumption, and oocyte growth, they may be enabled by a separate, lower minimal FSH threshold for a fraction of follicles. These results further supported the notion that FSH at 10 mIU/mL is near the upper bound of the FSH threshold range, which can drive nearly all follicles to mature, followed by fully-fledged oocyte maturation, ovulation, and luteinization. Furthermore, the suprathreshold levels of FSH of 20 or 30 mIU/mL significantly augment the secretion of preovulatory P4 and T without altering ovulatory events and post-ovulatory P4 secretion.

### FSH at different concentrations induces distinct expression patterns of follicle maturation-related genes

Follicles treated with various concentrations of FSH were collected on day 4 and 6 to examine the expression of genes crucial for follicle maturation using single-follicle RT-qPCR (Figure 3). The names, functions, and references of these genes are summarized in Suppl. Table 2 (33). Under 5 mIU/mL FSH, most genes stayed lowly expressed between day 4 and 6, including *Star*, *Cyp11a1*, *Cyp17a1*, *Cyp19a1*, *Lhcgr*, *Pappa*, *Inhba*, and *Nppc*, and the expression of *Hsd17b1* and *Fshr* was even significantly reduced on day 6. In contrast, FSH at 10 mIU/mL markedly induced the expression of most genes significantly, including *Cyp11a1*, *Hsd3b1*, *Hsd17b1*, *Cyp19a1*, *Lhcgr*, *Pappa*, *Inhba*, and *Nppc*. Compared with 10 mIU/mL, FSH at 20 and 30 mIU/mL further increased the expression of *Star*, *Cyp11a1*, and *Cyp17a1* but only statistically significant for *Cyp11a1*; the expression of other genes remained similar. *Amh*, a granulosa cell-specific gene highly expressed in small growing follicles, was reduced by 3.35-fold from day 4 to 6 when follicles were treated with 5 mIU/mL FSH, and was further diminished by 7.44-, 4.19-, and 7.61-fold by 10, 20, and 30 mIU/mL FSH, respectively. Collectively, these results are consistent with the phenotypical and hormonal data above, indicating that a minimal FSH threshold is required to induce the expression of follicle maturation-related genes. While the expression of several steroidogenic genes, such as *Star*, and *Cyp17a1*, and *Cyp11a1*, displayed a tendency for FSH concentration-dependent induction, other genes such as *Cyp19a1*, *Pappa*, and *Inhba*, which are primarily expressed in granulosa cells, are induced by FSH largely in an all-or-none manner.

**Figure 3.**
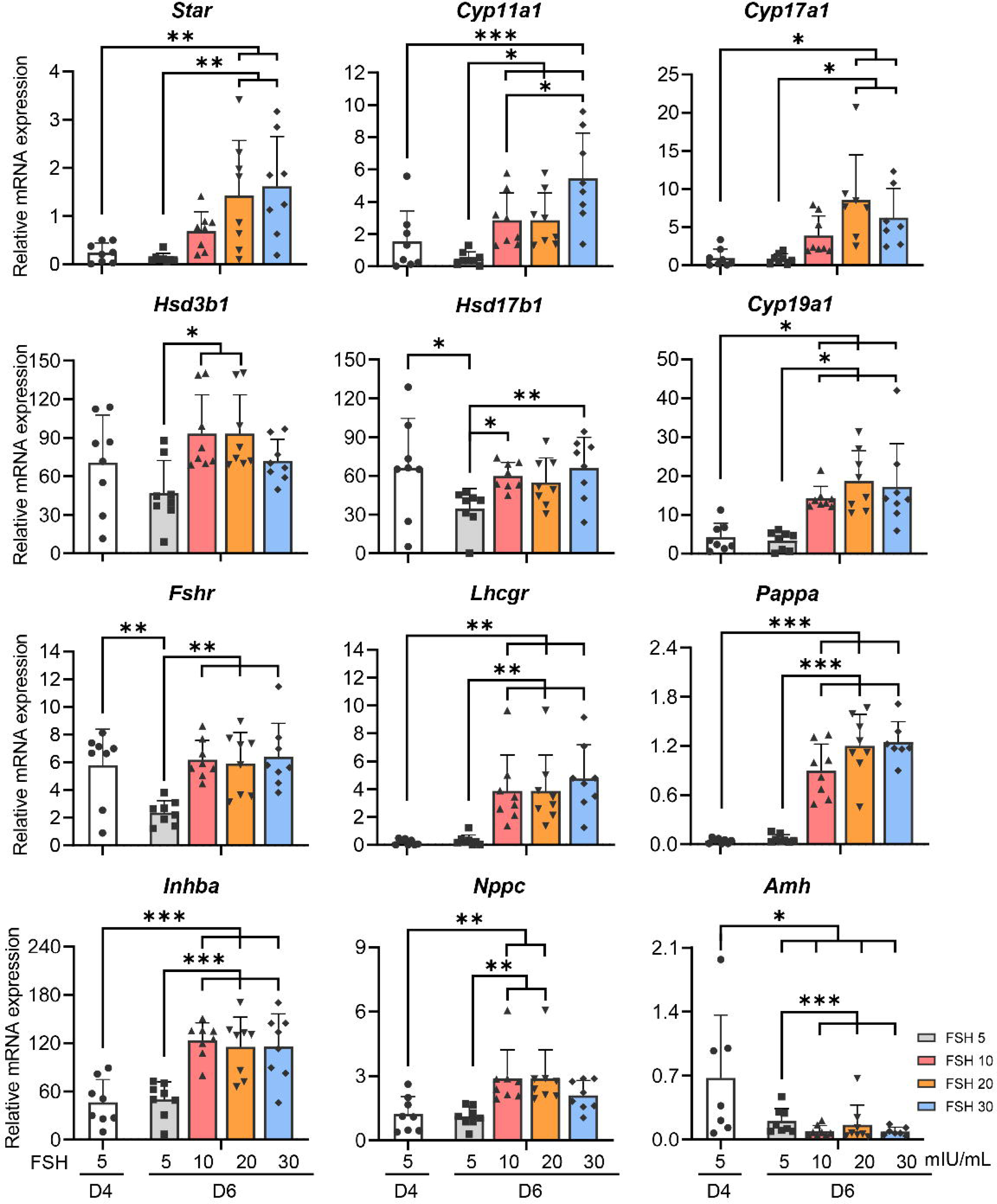
The concentration-response effects of FSH on the expression of follicle maturation-related genes in an *ex vivo* mouse folliculogenesis system. Follicles were cultured for 4 days with FSH of 5 mIU/mL to grow to the early antral stage, and then treated with FSH at 5, 10, 20 or 30 mIU/mL from day 4 to 6. Expression of steroidogenesis genes (*Star*, *Cyp11a1*, *Cyp17a1*, *Hsd3b1*, *Hsd17b1* and *Cyp19a1*) and follicle maturation-related genes (*Fshr*, *Lhcgr*, *Pappa*, *Inhba*, *Nppc* and *Amh*) were measured using RT-qPCR. N = 8 follicles in each group. Error bar: standard deviation; *\*P<0.05; ** P<0.01*.

### FSH at different concentrations induces distinct follicular transcriptomic profiles

To examine the follicular transcriptomics in a more unbiased manner, follicles cultured with various concentrations of FSH were collected at the end of day 6 for single-follicle RNA-seq analysis. PCA showed that follicles on day 4 formed a cluster that was clearly separated from follicles on day 6 cultured with 10-30 mIU/mL FSH, but they moderately overlapped with the day-6 follicles cultured with 5 mIU/mL FSH (Figure 4A). The latter formed an elongated cluster that bordered with the cluster of follicles cultured with 10 mIU/mL FSH. Follicles cultured with 20 and 30 mIU/mL FSH largely clustered together and tangibly overlapped with follicles cultured with 10 mIU/mL FSH. Collectively, these results indicate that follicles launch distinct transcriptomic responses to subthreshold through suprathreshold levels of FSH treatment, yet these responses of individual follicles collectively form a continuum in the transcriptomic space.

**Figure 4.**
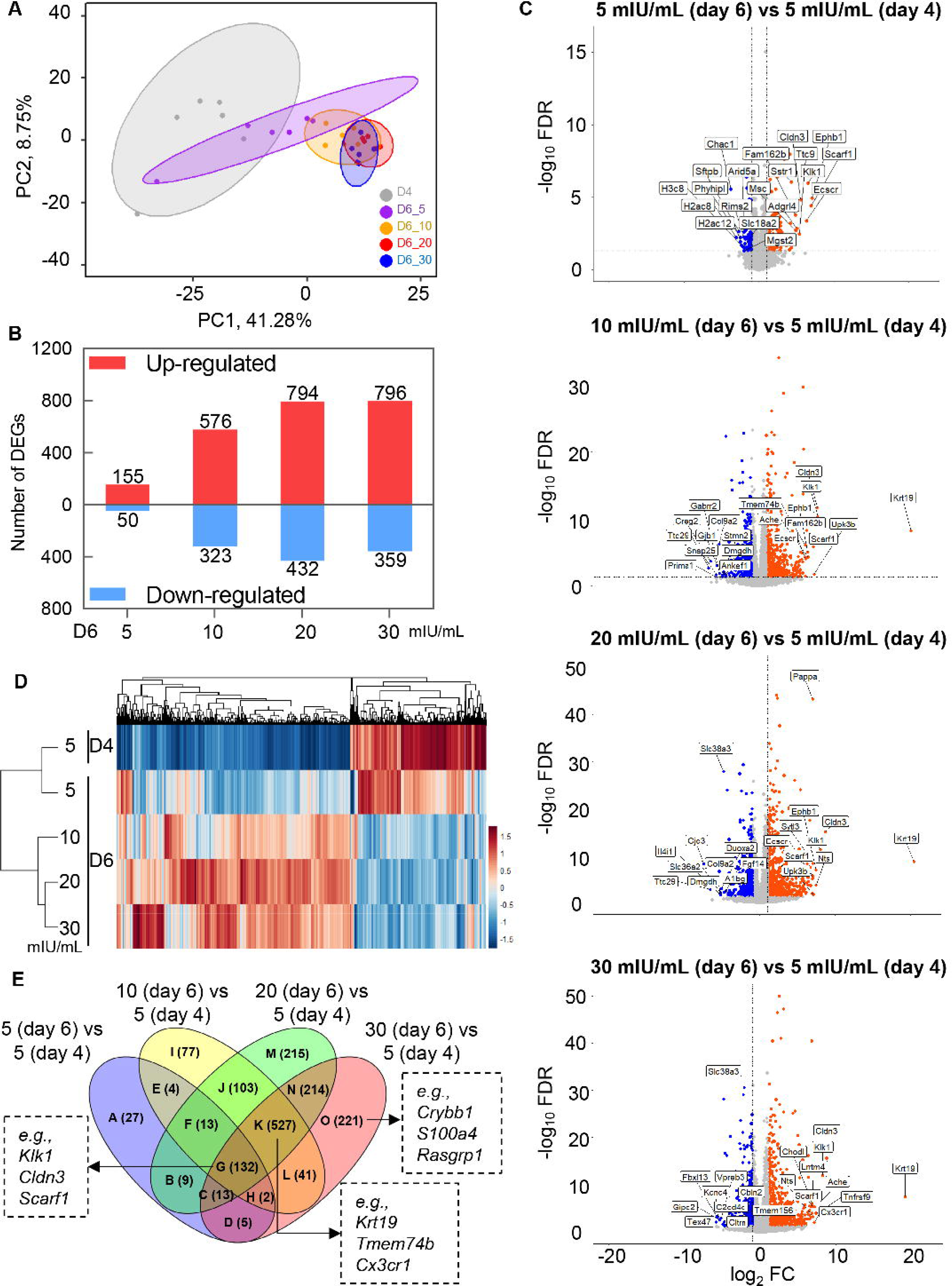
Concentration-response effects of FSH on follicular transcriptomics. (A) Principal component analysis (PCA) of the first two principal components among follicles on day 4 and follicles treated with different concentrations of FSH on day 6. (B) Number of DEGs expressed in follicles treated with FSH at 5, 10, 20 or 30 mIU/mL on day 6 compared to follicles on day 4. (C) Volcano plots of all genes, with top 10 genes identified. (D) Heatmap and hierarchical clustering based on the expression levels of DEGs in different treatment groups. (E) Venn diagram of DEGs. N = 6 follicles in each group.

DEGs were defined by a fold change of either > 2 or <0.5 and FDR < 0.05 between follicles from day 4 and follicles cultured with various FSH concentrations from day 4 to 6. Follicles treated with 5, 10, 20, and 30 mIU/mL FSH generated a total of 205, 899, 1226, and 1155 DEGs, respectively, among which 155, 576, 794, and 796 genes were up-regulated, and 50, 323, 432, and 359 genes were down-regulated (Figure 4B). All genes are listed in Suppl. Table 3 (33), and the top 10 up- and down-regulated DEGs are highlighted in the volcano plots in Figure 4C.

DEGs in all pair-wise comparisons were combined and used for hierarchical clustering analysis to further examine the transcriptomic relationships between follicles treated with different concentrations of FSH (Figure 4D). The heatmap showed that follicles were first clustered into two major groups at the top level. Group 1 were follicles from day 4 and follicles cultured with 5 mIU/mL FSH from day 4 to 6. Group 2 were follicles treated with 10-30 mIU/mL FSH. Within Group 2, Subgroup 2.1 included follicles treated with 10 mIU/mL FSH, while Subgroup 2.2 encompassed those treated with 20-30 mIU/mL FSH. In summary, the hierarchical clustering analysis is consistent with the PCA result and clearly demonstrates that 5 mIU/mL is a subthreshold FSH concentration that is insufficient to induce transcriptomic alteration characteristic of terminal follicle maturation; 10 mIU/mL FSH is close and slightly above the FSH threshold upper bound that is sufficient to induce the follicular transcriptomic changes required for follicle terminal maturation; and 20/30 mIU/mL FSH are suprathreshold concentrations that induce transcriptomic changes moderately similar to 10 mIU/mL FSH but also have a significant fraction of genes that are differentially expressed.

The Venn plot in Figure 4E summarized the numbers of overlapping and unique DEGs between FSH pair-wise comparisons. Each section represents genes specifically or commonly altered by one or multiple FSH concentrations. For instance, section G consisted of 132 genes consistently regulated from day 4 to 6 by all FSH concentrations (e.g., *Klk1*, *Cldn3*, and *Scarf1*); the genes in sections I (77), J (103), K (527), and L (41) combined were selectively altered when FSH increased from 5 to 10 mIU/mL; the 527 genes in section K were commonly up- or down-regulated in 10, 20, and 30 FSH groups (e.g., *Krt19*, *Tmem74b*, and *Cx3cr1*). In section O, there were 221 genes selectively changed by the suprathreshold level of 30 mIU/mL FSH (e.g., *Crybb1*, *S100a4*, and *Rasgrp1*). Detailed gene lists for each section are provided in Suppl. Table 4 (33).

### FSH at different concentrations alters distinct gene regulatory pathways

DEGs in all pair-wise comparisons in Figure 4B were used for the GO and KEGG pathway analysis. All significantly enriched GO terms and KEGG pathways are summarized in Suppl. Table 5 (33), with the top 10 highlighted in Figure 5. There were common and unique GO terms in different FSH concentration comparisons. For instance, 78 GO terms were commonly enriched in all four FSH comparisons, including 38 biological process (BP), 27 cellular component (CC), and 13 molecular function (MF) (Suppl. Figure 1) (33). These commonly altered GO terms are related to the BP of ‘angiogenesis’, ‘cell adhesion and morphogenesis’, and ‘neural processes’, etc.; CC of ‘extracellular components’, ‘membrane components’, and ‘cellular junctions’, etc.; and MF of binding activities, such as ‘insulin-like growth factor binding’. Compared to 5 mIU/mL FSH, more GO terms (n=277) were selectively enriched in the 10, 20, and 30 mIU/mL FSH groups, including 182 BP related to ‘cell adhesion’, ‘migration and development’, cellular responses and regulation such as ‘response to estradiol’, regulation of biological processes such as ‘cAMP-mediated signaling’, 48 CC associated with ‘plasma membrane and junctions’ and ‘cytoplasmic structures’, etc., and 47 MF linked to binding activities such as ‘extracellular matrix binding’ and enzyme activities such as ‘metalloendopeptidase activity’. Compared to 5 and 10 mIU/mL, there were 85 enriched GO terms in FSH of 20 and 30 mIU/mL, including 56 BP, 11 CC, and 18 MF. These GO terms were primarily related to the BP of ‘cell adhesion’, ‘migration and development’, ‘cell signaling and regulation’, and other processes such as ‘regulation of oocyte maturation’; CC of ‘synaptic components’ and ‘cell junctions’, etc.; and MF of binding activities such as ‘Wnt-protein binding’ and enzyme activities such as ‘cytokine binding’.

**Figure 5.**
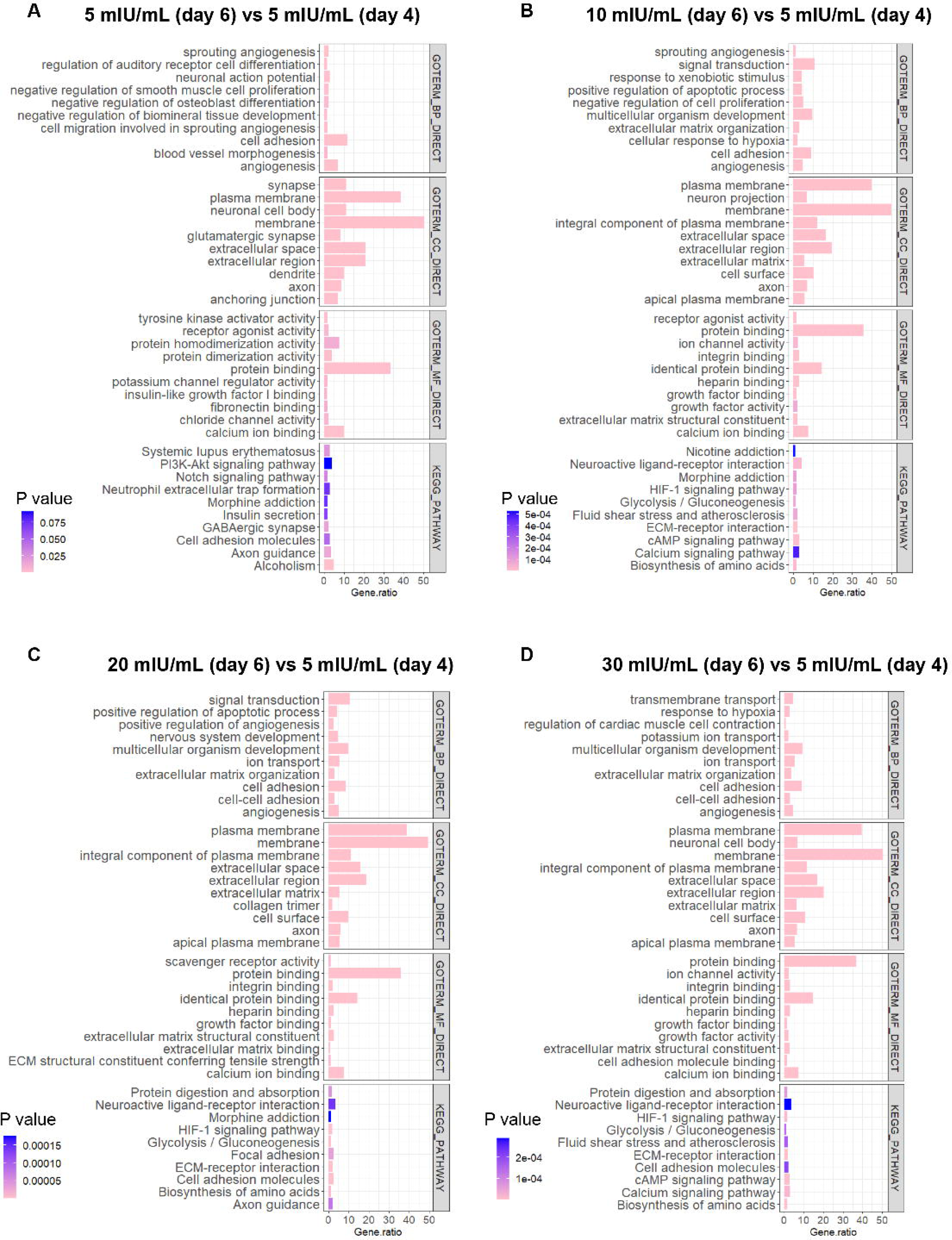
Enriched Gene Ontology (GO) and Kyoto Encyclopedia of Genes and Genomes (KEGG) pathways associated with DEGs in follicles treated with FSH at 5, 10, 20 or 30 mIU/mL on day 6 compared to follicles on day 4. The top 10 GO biological process (BP), molecular function (MF) and cellular component (CC) and top 10 KEGG pathways are shown.

Regarding the significantly enriched KEGG pathways, three of them were altered by all four FSH concentrations, including ‘GABAergic synapse’, ‘Cell adhesion molecules’, and ‘Axon guidance’ (Suppl. Figure 2A) (33). There were also KEGG pathways selectively altered by a specific FSH concentration. Compared to 5 mIU/mL FSH, 29 pathways (Suppl. Figure 2B) (33) were selectively changed by 10-30 mIU/mL FSH, such as ‘Fluid shear stress and atherosclerosis’, ‘ECM-receptor interaction’, ‘cAMP signaling pathway’, ‘HIF-1 signaling pathway’, ‘MAPK signaling pathway’, and “Ras signaling pathway”. Notably, several of these 29 KEGG pathways were related to the metabolism, including ‘Metabolic pathways’, ‘Central carbon metabolism in cancer’, ‘Cortisol synthesis and secretion’, ‘Biosynthesis of amino acids’, ‘Carbon metabolism’, ‘Starch and sucrose metabolism’, ‘Glycine, serine and threonine metabolism’, ‘Arginine biosynthesis’, and ‘Glycolysis/Gluconeogenesis’. For all enriched metabolic pathways, we used involved DEGs to perform a similar hierarchical clustering analysis as we performed in Figure 4D. The results showed that the expression pattern of these metabolism-related DEGs was FSH concentration-dependent (Figure 6A). Compared to follicle from day 4 or follicles treated with 5 mIU/mL FSH from day 4 to 6, 10-30 mIU/mL FSH induced similar transcription patterns of these genes. The interaction of multiple metabolic genes (Figure 6B) further revealed the functional synergy among different metabolic processes, and the top-ranking genes of each process were identified (orange nodes) via Cytohubba. *Tpi1*, *Pkm*, and *Pgk1* shared by biosynthesis of amino acids, carbon metabolism, and glycolysis exhibited in a similar expression pattern, suggesting a high degree of coregulation (Figure 6C). For the Glycine, serine, and threonine metabolism, the top-ranking genes were down-regulated by FSH at 5-30 mIU/mL. For glycolysis, *Aldob*, *Aodoc*, and *Eno2* were concentration-dependently up-regulated. For Starch and sucrose metabolism, the expression of *Pygl* and *Pgm1* was increased, but *Pygb* was reduced by FSH from 5 to 30 mIU/mL. For Arginine biosynthesis, the expression of *Ass1* was obviously dependent on FSH concentrations. For Cortisol synthesis and secretion, the top-ranking genes were up-regulated in a concentration-dependent manner. In contrast to 5-10 mIU/mL FSH, 3 KEGG pathways were selectively altered by 20 or 30 mIU/mL FSH, including ‘Amoebiasis’, ‘Malaria’, and ‘Complement and coagulation cascades’, which were related to immune response (Suppl. Figure 2C) (33). These findings suggest a dose-dependent effect of FSH at the regulatory pathway level, highlighting a crucial role of metabolism-related pathways during FSH-induced follicle maturation and related follicular functions.

**Figure 6.**
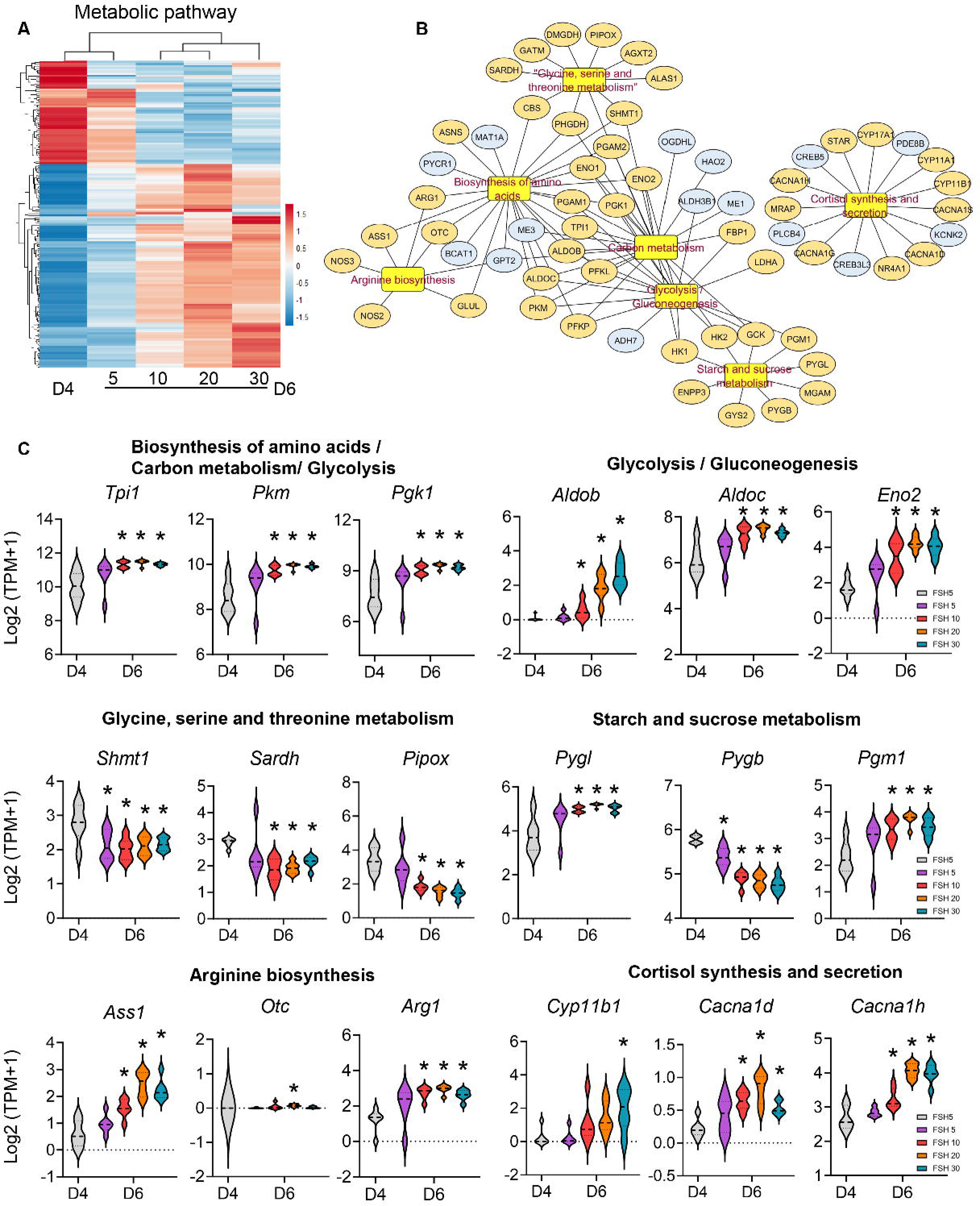
Concentration-response effects of FSH on follicular metabolic pathways in an *ex vivo* folliculogenesis system. (A) Heatmap of genes in the metabolic related pathways. (B) The interaction of metabolic pathway related genes. Top three ranking genes were labeled in yellow. (C) mRNA expression levels (log2[TPM + 1]) of metabolic related genes in follicles on day 4 and on day 6 after treatment of FSH at 5, 10, 20 and 30 mIU/mL.

### Excessive FSH induces proinflammatory responses in follicular cells

Since the GO and KEGG analysis revealed changes of multiple gene regulatory pathways when follicles were treated with suprathreshold concentrations (20 and 30 mIU/mL) of FSH, we next performed a more in-depth comparison of the follicular transcriptomics between the 10 and 30 mIU/mL FSH groups to gain further insights into the molecular mechanisms underpinning the potential detrimental effect of follicle stimulation by excessive FSH. Using the same criterion as shown in Figure 4, there were 60 DEGs, with 58 genes up-regulated and 2 genes down-regulated in follicles treated with 30 mIU/mL FSH. All genes are summarized in Suppl. Table 3 (33). The top 10 up-regulated and 2 down-regulated DEGs are highlighted in the volcano plot in Figure 7A, and the top 10 enriched GO terms and KEGG pathways were presented in Figure 7B. FSH at 30 mIU/mL altered several GO terms and KEGG pathways crucial for FSH-dependent follicle maturation (39–41), including ‘G protein-coupled receptor signaling pathway’, ‘cellular response to cAMP’, ‘glucocorticoid receptor binding’, ‘PI3K-AKT signaling pathway’, and ‘ovarian steroidogenesis’, suggesting that suprathreshold FSH further induced signaling molecules and expression of FSH target genes to higher levels.

**Figure 7.**
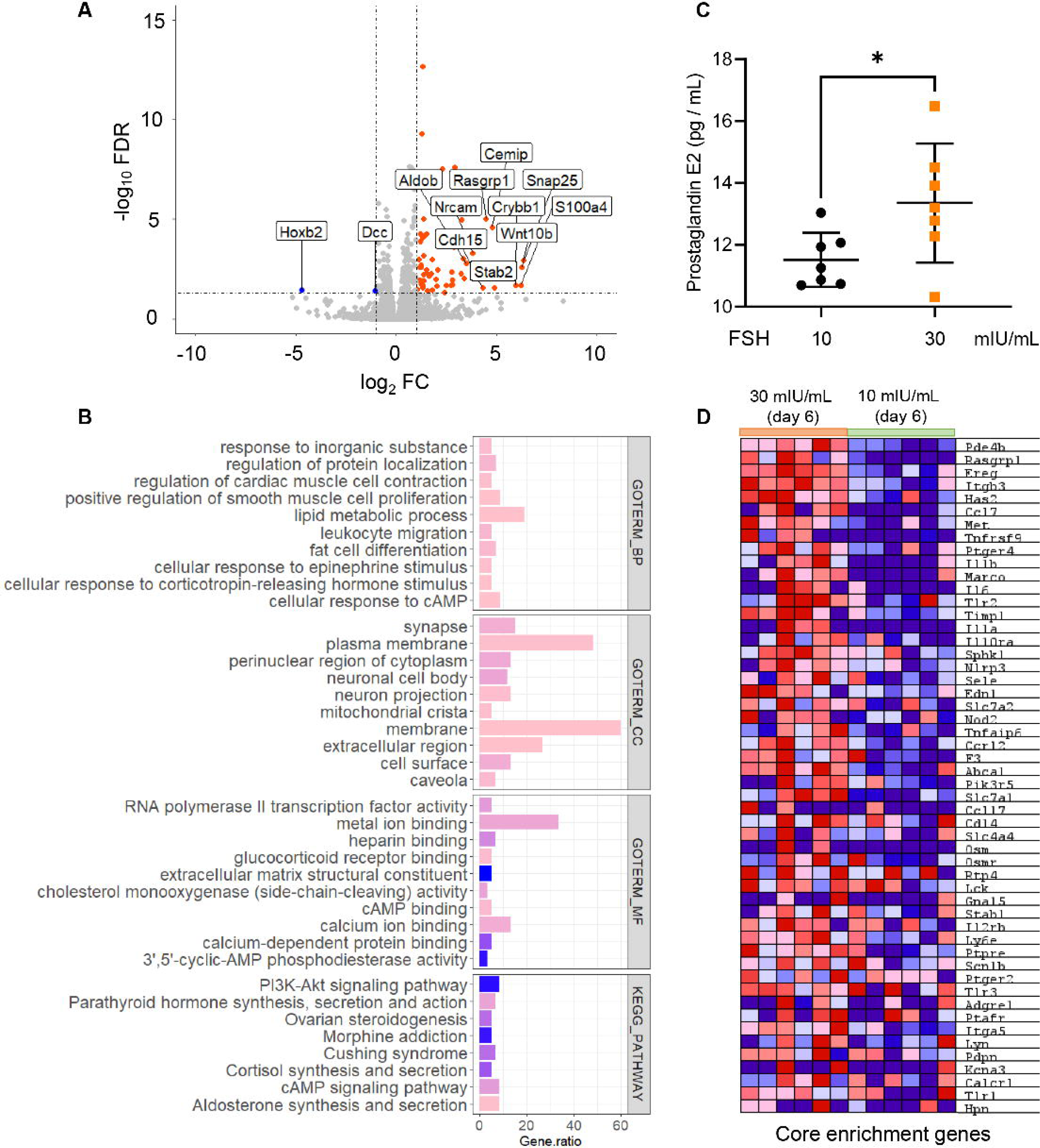
The transcriptome-wide changes of genes and signaling pathways in follicles treated with the excessive FSH of 30 mIU/mL compared with 10 mIU/mL on day 6. (A) Volcano plot of all genes, with the top 10 up- and down-regulated genes highlighted. (B) Top 10 GO enrichment of BP, MF and CC and top 10 KEGG pathway enrichment of DEGs expressed in follicles exposed to FSH of 30 mIU/mL compared to 10 mIU/mL. (C) Concentration of PGE2 in the conditioned follicle culture media on day 6 (N=7). (D) Heatmap of genes in the inflammatory response pathway. Error bar: standard deviation; *\*P<0.05*.

Among all DEGs, it is intriguing to note that there were many established ovulation or luteinization-related genes up-regulated by 30 mIU/mL FSH, including *Sfrp4* (9.5-fold), *Ptgs2* (7.45-fold), *Ptgfr* (6.77-fold), *Star* (2.91-fold), *Ereg* (2.53-fold), *Runx2* (2.35-fold), and *Cyp11a1* (2.24-fold). PTGS2, also termed cyclooxygenase-2 or COX-2, is crucial for ovulation by promoting PGE2 synthesis related inflammatory responses (42). Consistent with the transcriptional induction of *Ptgs2*, the ELISA data confirmed that at the hormonal level, follicles treated with 30 mIU/mL FSH had significantly increased secretion of PGE2 compared to 10 mIU/mL FSH (Figure 7C). Gene Set Enrichment analysis (GSEA) further revealed the significant enrichment of inflammatory signaling pathways in follicles treated with 30 mIU/mL FSH, with the enrichment score of 0.5 and *p* < 0.05 (Figure 7D). The heatmap in Figure 7D showed that the core enrichment of inflammatory genes and their expression levels in follicles treated with 30 mIU/mL FSH were significantly different from follicles treated with 10 mIU/mL FSH, including genes encoding cytokines (*Il6*, *Il1a*, *Il1b*, and *Tnfaip6*), chemokines (*Ccl7* and *Ccl17*), and cytokine receptors (*Cxcr4, Ccrl2,* and *Tnfrsf9*). These results suggest that there are proinflammatory responses in follicles treated with a suprathreshold concentration of FSH.

### Soft clustering analysis identifies distinct concentration-response relationship of FSH target genes and underlying molecular events

We next used DEGs combined from all pairwise comparisons between day 4 and various FSH concentration groups on day 6 for the M-fuzzy clustering analysis to determine the concentration-response relationship of FSH on follicular transcriptomics. Six clusters were enriched, with specific genes in each cluster summarized in Suppl. Table 6 (33). Overall, DEGs exhibited either monotonically increasing (Cluster 1 and 2), decreasing (Cluster 4 and 5), or nonmonotonic (Cluster 3 and 6) expression patterns (Figure 8).

**Figure 8.**
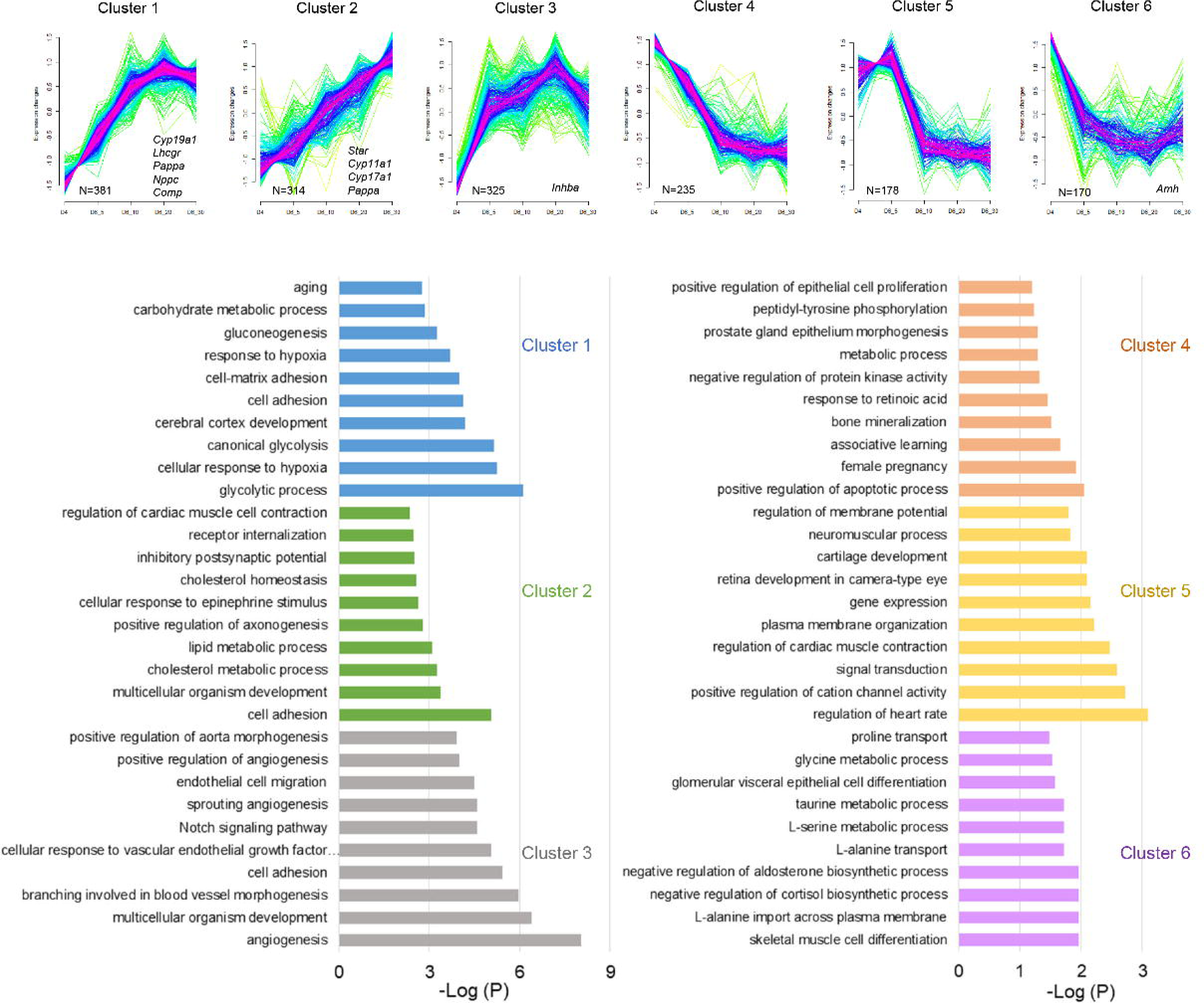
Soft clustering analysis of DEGs in follicles on day 4 and day 6 treated with 5, 10, 20 and 30 mIU/mL. GO analysis of DEGs in each identified cluster, with top 10 enriched biological process terms of each cluster shown.

The expression levels of genes in Cluster 1 continuously increased when follicles were treated with FSH up to 10 mIU/mL, but 20 and 30 mIU/mL FSH only slightly increased their expression. Several well-established follicle maturation-related genes expressed in the granulosa cells were primarily enriched in Cluster 1, including *Cyp19a1*, *Lhcgr*, *Nppc*, and *Comp*. The genes in Cluster 2 exhibited a more apparent concentration-dependent increase pattern, including *Pappa*, *Cyp11a1*, and theca cell-specific genes *Star* and *Cyp17a1*. The genes in Cluster 4 continuously decreased but the higher levels of FSH did not result in a further reduction. For genes in Cluster 5, there was no significant change from day 4 to 6 until follicles were treated with 10 mIU/mL FSH or above, indicating that a minimum FSH is required for their transcriptional repression. Genes in cluster 3 and 6 remarkably increased or decreased from day 4 to 6 at all FSH concentrations, suggesting that the expression of these genes are more sensitive to the time or temporal change of *ex vivo* folliculogenesis; however, their expression also reached a peak or nadir at 20 mIU/mL but reversed the direction at 30 mIU/mL FSH.

We next used DEGs in each cluster with a membership cutoff of > 0.6 for the GO BP analysis to gain more mechanistic insights (Figure 8). Genes in Cluster 1 were primarily related to ‘glycolytic process’, ‘adhesion and hypoxia’. Genes in Cluster 2 were associated with ‘adhesion’, ‘cholesterol’, and ‘lipid metabolism’. Genes in Cluster 3 were related to ‘angiogenesis’ and ‘cell adhesion’. Genes in Cluster 4 were linked to ‘regulation of apoptosis’, ‘pregnancy’, and ‘protein kinase activity’. Genes in Cluster 5 were related to ‘regulation of cation channel activity’ and ‘membrane organization’. Genes in Cluster 6 were associated with ‘negative regulation of cortisol’ and ‘aldosterone biosynthesis’, ‘L-alanine transport’, and ‘amino acid (L-serine, taurine, and glycine) metabolic process’.

### Excessive FSH does not alter ovulatory genes or genes related to inflammation and vascular permeability in individual follicles

Women treated with high doses of FSH for ovarian stimulation are at an increased risk of OHSS which has been related to hCG-induced overstimulation of molecules related to inflammation and vascular permeability (43,44). Thus, despite that our results in Figure 2G-2H showed no significant changes in ovulation and luteinization outcomes when follicles were treated with the suprathreshold levels of 20-30 mIU/mL FSH, we further examined whether excessive FSH sensitizes the expression of ovulation-related genes, particularly genes associated with angiogenesis and inflammation, in response to the hCG stimulation for ovulation induction. Follicles treated with 10 or 30 mIU/mL FSH from day 4 to 6 were collected at 0 or 4-hour post-hCG for single-follicle RT-qPCR to examine the expression of key ovulatory genes. The expression of most LH/hCG target genes in follicle has been observed to peak at 4 hours (45). The specific gene names and functions are summarized in Suppl. Table 7 (33). Similar to the results in Figure 3, at 0 hr, follicles treated with 30 mIU/mL FSH exhibited significantly higher expression levels of follicle maturation and steroidogenic genes, including *Lhcgr*, *Cyp19a1*, *Star*, and *Cyp11a1* (Figure 9). Follicles treated with 30 mIU/mL FSH also had higher expression levels of three ovulatory genes, *Ereg*, *Ptgs2*, and *Runx2* before the hCG treatment. At 4 hours after hCG treatment, most ovulation and luteinization-related genes, such as genes related to vascular permeability and cumulus cell expansion, had comparable expression levels between 10 and 30 mIU/mL-treated follicles except that *Plau* was slightly decreased. These results suggest that the excessive suprathreshold level of FSH does not further promote the expression of ovulatory genes in individual follicles.

**Figure 9.**
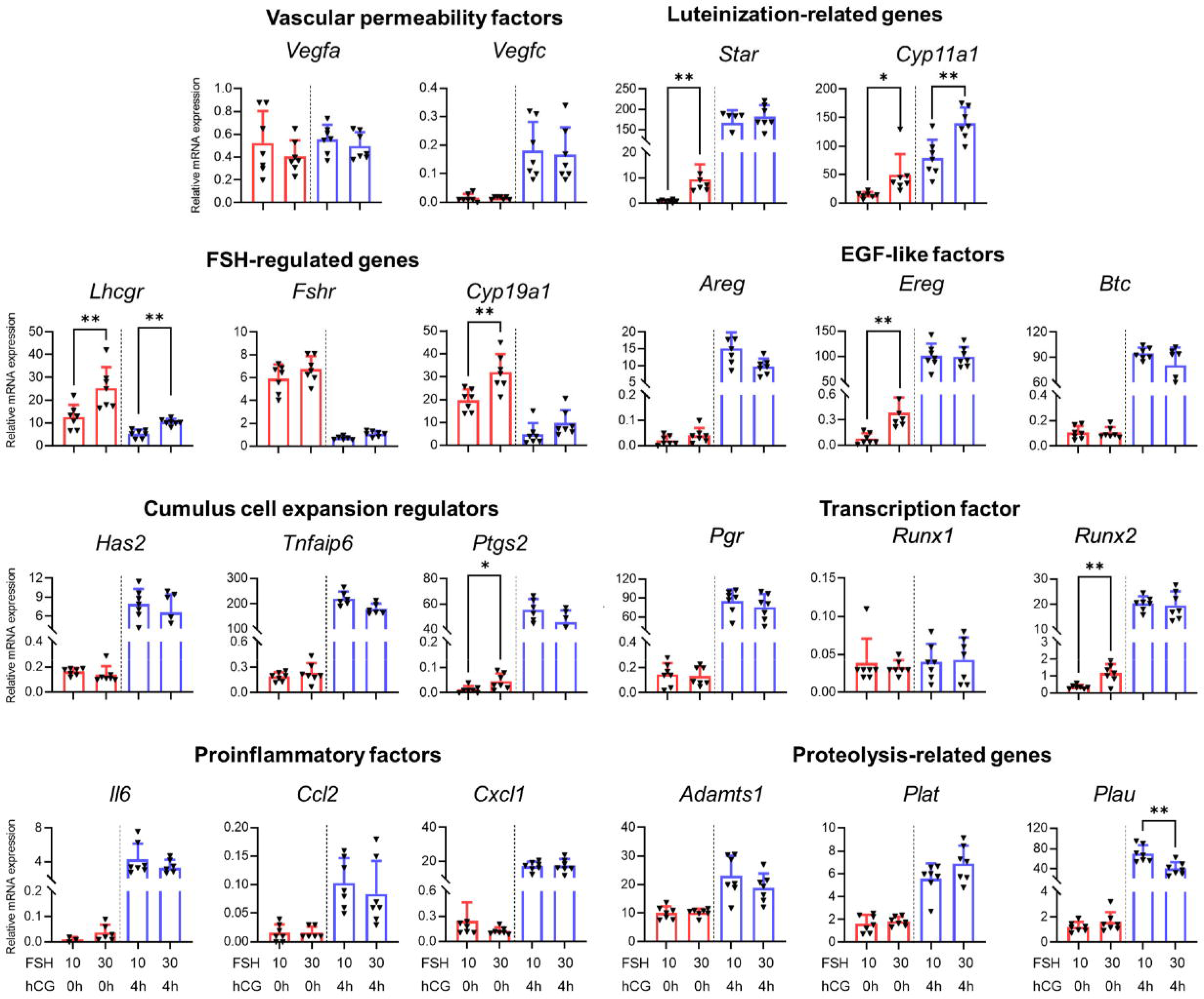
mRNA expression levels of ovulatory genes examined by RT-qPCR in follicles treated with 10 and 30 mIU/mL at 0 and 4 hours post-hCG. N = 7 follicles in each group. *\*P<0.05; ** P<0.01*.

### Excessive FSH does not affect oogenic expression of maternal effect genes but reduces the expression of genes related to mitochondria and energy metabolism

In parallel to folliculogenesis, the follicle-enclosed oocyte produces and accumulates maternal mRNAs, proteins, and organelles to acquire both meiotic and developmental competence, which are essential for subsequent fertilization, early embryogenesis, and zygotic genome activation (12). Our results in Figure 2 showed that 20-30 mIU/mL FSH did not further alter oocyte size and resumption and completion of meiosis I compared with 10 mIU/mL. Here, we further examined whether excessive FSH affects the oocyte transcriptome. Follicles were treated with FSH at 10 or 30 mIU/mL from day 4-6 in eIVFG, and ovulated MII oocytes were collected at 14 hours post-hCG for single-oocyte SMART-Seq2 RNA-seq analysis. All genes in oocytes are summarized in Suppl. Table 8. PCA showed that the oocytes from follicles treated with FSH at 10 and 30 mIU/mL vastly overlapped, and there was a higher degree of consistency among oocytes in the 30 mIU/mL FSH group (Figure 10A). Pearson’s correlation analysis revealed that the oocyte transcriptomic profiles between 10 and 30 mIU/mL FSH groups were highly similar with a correlation coefficient (R) of 0.99167 (Figure 10B, left panel). When we compared several oocyte marker genes, including *Gdf9*, *Bmp15*, and *Zp2*, their expression levels were comparable between the two FSH groups (Figure 10B, right panels). We next examined a list of 70 identified maternal effect genes (Suppl. Table 9) (12), such as *Cdc20*, *Ooep*, and *Padi6*, which also showed nearly identical expression levels between the two FSH groups, with a high Pearson’s correlation coefficient (R) of 0.99483 (Figure 10C).

**Figure 10.**
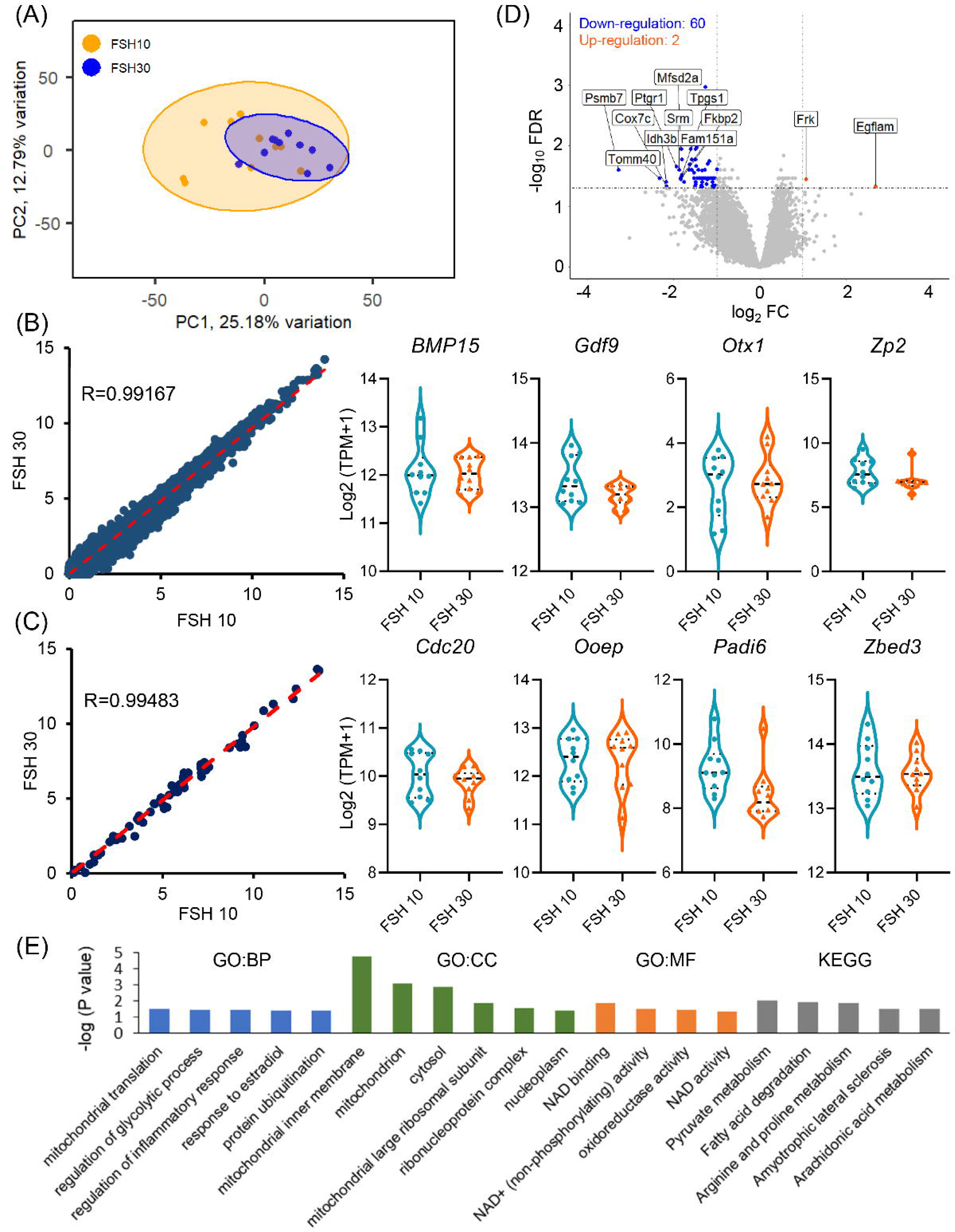
Effects of threshold and suprathreshold levels of FSH on oocyte transcriptomics. (A) PCA of the first two principal components among oocytes from follicles treated with FSH of 10 and 30 mIU/mL. (B) Pearson’s correlation analysis of gene expression in oocytes from follicles treated with 10 and 30 mIU/mL. mRNA expression levels (Log2 (TPM+1)) of representative oocyte makers. (C) Pearson’s correlation analysis of expression of maternal effect genes in oocytes from follicles treated with FSH of 10 and 30 mIU/mL. (D) Volcano plot of all genes in oocytes, and 2 significantly up-regulated genes and top 10 down-regulated genes were highlighted. (E) Top 5 GO terms of BP, MF and CC and top 5 KEGG pathways of DEGs in oocytes from follicles treated with FSH at 10 and 30 mIU/mL on day 6.

Using the same criterion identifying DEGs as shown in Figure 4, there were only 62 DEGs between two FSH concentration groups, with 2 genes up-regulated and 60 genes down-regulated in oocytes from follicles treated with excessive 30 mIU/mL FSH (Figure 10D). So far, most DEGs have not been reported to essentially regulate oogenesis. There were 20 significantly enriched GO terms and KEGG pathways based on the 62 DEGs, and most of them were related to mitochondria and energy metabolism, such as ‘mitochondrial translation’, ‘NAD activity’, and ’pyruvate metabolism’ (Figure 10E). In summary, these results demonstrate that a suprathreshold level of FSH does not appear to alter the overall oocyte transcriptomics or the expression of specific oocyte marker genes or maternal effect genes; however, some transcripts related to oocyte maturation, fertilization, mitochondria, and energy metabolism were altered.

## Discussion

Gonadotropin-dependent folliculogenesis, a crucial step for supporting the eventual follicle maturation, ovulation, luteinization, and reproductive cycles and fertility, is well conserved across many mammalian species. Both insufficient and excessive FSH have been linked to poor female reproductive outcomes (1,2,21–25,29,30). Here we performed a dose-response functional and transcriptomic analysis of FSH using an *ex vivo* folliculogenesis and oogenesis system. Our results demonstrate that (1) a minimum FSH threshold is required to support follicle maturation and related ovarian functions, and such threshold is moderately variable among individual follicles; (2) excessive or suprathreshold levels of FSH augment preovulatory secretion of P4 and T; (3) FSH at the subthreshold, threshold, and excessive levels induces distinct expression patterns of follicle maturation-related genes and distinctly altered the entire follicular transcriptomics; (4) excessive FSH promotes proinflammatory responses in follicular cells; (5) excessive FSH does not alter ovulation and OHSS-related genes in individual follicles; and (6) excessive FSH does not alter maternal effect genes but reduces the expression of genes related to mitochondria and energy metabolism in oocytes.

Although the concept of FSH threshold is invoked primarily for mono-ovulatory species in the literature, it is highly likely that in poly-ovulatory species FSH also has to rise to certain levels to stimulate the eventual maturation of multiple antral follicles toward the preovulatory stage. As demonstrated with the mouse follicles cultured *ex vivo* here, each follicle has its own FSH threshold to overcome in order to be stimulated into a discrete, high E2-secreting state with a characteristic transcriptomic profile (Figure 1D and 2D). The range of the FSH threshold seems between 5-10 mIU/mL for the mouse follicles *ex vivo*, below which no follicles and above which nearly all follicles mature into the high E2-secreting state with basically all-or-none induction of certain genes including *Cyp19a1* (Figure 1-3). It is likely that in mice *in vivo* the physiological FSH can rise past the thresholds of a number of follicles, stimulating them to mature into the preovulatory stage. Although differential threshold was demonstrated with mouse follicles here, it is in keeping with the mono-ovulatory species, for which among the cohort of recruited antral follicles only the most sensitive one, i.e., the follicle at the most advanced stage or has the most expression levels of FSHR in granulosa cells and thus expected to need the lowest FSH to become dominant.

Interestingly the FSH threshold does not seem to apply to all the phenotypical events of terminal follicle maturation. Although follicle growth, as measured by follicle diameters, is FSH concentration-dependent, even at the lower bound of the FSH threshold range, i.e., 5 mIU/ml, the follicles still seem to grow at a robust rate (Figure 1-2). Our single-follicle RNA-Seq analysis also revealed the comparable expression levels of cell-cycle related genes, such as *Ccna2*, *Ccnb1*, *Ccne2*, and *Cdc25a*, in different concentrations of FSH at 5-30 mIU/mL (Suppl. Table 3). Similarly, over half of the follicles can manage to ovulate with high oocyte meiosis resumption at 5 mIU/mL FSH (Figure 2G), despite that there is barely any *Lhcgr* gene induction at the transcript level (Figure 3). Given the multitude of signaling pathways triggered by FSH (19,46–48), it is possible that different maturation events, such as granulosa cell proliferation, E2 secretion, and preparation for ovulation, are underpinned by different pathways, thus requiring different levels of FSH stimulation. In our study, excessive FSH at > 10 mIU/mL slightly but not significantly promoted follicle growth, aromatase induction, E2 secretion, and ovulation rate (Figure 1-3). Similarly, previous studies also found that the increase of E2 secretion start to level off with FSH at 100 ng/mL and 10 mIU/mL in KGN cell model and a two-layered mouse follicle model (49,50). These results indicated that there is an all-or-none response where additional FSH does not result in significant increase in induction of aromatase activity.

Premature luteinization is defined as an early elevation of circulating P4 in the late follicular phase, measured prior to or on the day of hCG trigger in women receiving ovarian stimulation for IVF (51,52). It often causes a cancellation of fresh embryo transfer due to the risks of altered endometrial receptivity and embryo-endometrial asynchrony, which result in lower implantation and pregnancy success (52–56). High doses of FSH have been postulated as a major reason for the high P4 secretion (57), but the underlying mechanisms remain unknown. In two previous studies evaluating 110 and 81 women undergoing ovarian stimulation, 13.4% and 38.8% of IVF cycles experienced premature luteinization, respectively (54,57). Similar to these *in vivo* studies, in our *ex vivo* system we found that excessive FSH significantly increased pre-ovulatory P4 secretion (Figure 2E). At molecular levels, it promoted the transcription of steroidogenic genes of *Star*, *Cyp17a1*, and *Cyp11a1* (Figure 3), all of which critically contribute to P4 synthesis. However, excessive FSH did not increase the expression of *Hsd3b1* (Figure 3), a gene encoding the hydroxylase enzyme for converting pregnenolone to P4. These results suggest that ovarian stimulation-induced premature luteinization may be primarily caused by the transcriptional induction of *Star*, *Cyp17a1*, and *Cyp11a1*, highlighting potential adjuvant therapeutic targets to prevent premature luteinization during ovarian stimulation for IVF. It is unknown how excessive FSH promotes the transcription of theca cell-specific genes of *Star* and *Cyp17a1*, as FSHR is expressed in granulosa cells. We speculate that ligands induced by excessive FSH in granulosa cells, such as IGF, may act as paracrine factors on theca cells to promote the expression of theca cell-specific steroidogenic genes(17,19), which eventually result in a high P4 secretion and premature luteinization.

Our ELISA data demonstrated a notable increase of preovulatory T secretion in follicles treated with excessive FSH (Figure 2F). There is currently limited evidence regarding the effects of ovarian stimulation on androgen production. During IVF treatment, elevated T levels have been observed in several patients with ovarian hyperstimulation syndrome or pretreatment of GnRH agonist (58,59). Another study focusing on a single patient with isolated FSH deficiency, reported that the administration of combined FSH and hCG caused up to 2-fold increase of circulating T and its precursor of androstenedione compared to the hCG treatment alone (60). PCOS patients treated with high levels of FSH have also been found to have increased circulating levels of 17-hydroxyprogesterone, androstenedione, dehydroepiandrosterone (61). The potential impacts of hyperandrogenism in women receiving high doses of FSH may be due to the induction of CYP17A1 in theca cells and FSH-induced inhibin and IGF in granulosa cells (62,63). It has been demonstrated in theca cells of many species that androgen production can be increased by inhibin (64–68). IGF-1 can also stimulate basal and potentiate LH-stimulated androgen production in both human and rat thecal cells (66,69,70). Our RT-qPCR (Figure 3) and RNA-seq data (Suppl. Table 3) showed that the expression of all inhibin-related genes, including *Inhba*, *Inhba*, and *Inhbb*, remained however similar in the 10-30 mIU/mL FSH groups. We hypothesize that the increased secretion of T in our *ex vivo* system may be just secondary to the high P4 production in excessive FSH-treated follicles, which eventually results in an elevated T synthesis. High T secretion has been shown to impair early embryo development and uterine receptivity in mice, leading to delayed or failed embryo implantation (71). In summary, these results indicate that women receiving high levels of FSH administration may experience hyperandrogenism and associated impairment of early pregnancy events of early embryogenesis, uterine receptivity, and implantation.

Compared to naturally cycling women, women receiving ovarian stimulation for infertility treatment have been shown to have higher expression levels of proinflammatory genes in granulosa cells of preovulatory follicles as well as higher secretion levels of cytokines in the follicular fluid, including IL6, IL10, IL18, and TNF (72,73). The GSEA result using our single-follicle RNAs-seq data revealed that excessive FSH significantly promoted the transcription of a number of genes encoding proinflammatory factors, including cytokines (e.g., *Il1a*, *Il1b*, and *Tnfaip6*), *Ptgs1*, *Ptgs2*, and several ovulatory genes (e.g., *Ereg* and *Runx2*) (Figure 7D). Ovulation has been well established as an inflammatory process (74). While these proinflammatory factors are essential for LH/hCG-induced ovulation, the early induction of these genes in preovulatory follicles appear to be insufficient to trigger ovulation but may cause a premature influx of leukocytes, potentially impairing the FSH-stimulated terminal follicle maturation and subsequent ovulation (75,76). PGE2, a potent proinflammatory mediator, is synthesized through the metabolism of arachidonic acid by PTGS2 (77). Our results showed that both the transcriptional induction of *Ptgs2* and the hormonal secretion of PGE2 were highly induced by the excessive FSH (Figure 7). Collectively, these results highlight a potential risk of ovarian inflammation induced by the administration of high doses of FSH.

OHSS is a potential complication of ART, particularly in individuals with PCOS or those who receive high doses of FSH for ovarian stimulation followed by a hCG trigger for the induction of oocyte maturation (78). OHSS is characterized by enlarged ovarian cysts and seeping of intravascular fluids into abdominal or pleural compartments (79). So far, the pathophysiology of OHSS remains poorly understood, but it has been related to elevated circulating levels of E2 during ovarian stimulation and the induction of vascular endothelial growth factors (VEGFs) in granulosa cells following hCG stimulation (80), which cause increased capillary permeability in the ovary and/or systemic circulation. Our results showed that compared to 10 mIU/mL FSH, excessive FSH at 30 mIU/mL did not alter the expression of VEGF-related genes in individual follicles before or after hCG treatment (Figure 9), suggesting that follicular VEGFs may not be regulated by excessive FSH, at least at the single follicular level. Rather, systemic VEGF levels may depend on the number of recruited large antral follicles. This notion is consistent with the findings in several previous clinical studies. For instance, a high risk of OHSS only occurred in IVF patients with more than 11 antral follicles of >= 10 mm in diameter on the day of hCG trigger (81), and lowering FSH reduced the numbers of preovulatory follicles as well as lowered the risk of OHSS (82,83). These results suggest that with the administration of high doses of FSH, it is primarily the induction of more preovulatory follicles rather than the sensitization of individual follicles to hCG stimulation to secrete more VEGFs that leads to higher systemic levels of VEGFs, increased vascular permeability, and heightened risk of OHSS. In addition to VEGFs, the transcriptional induction of *Lhcgr*, *Ptgs2*, and several proinflammatory cytokines (e.g., IL-1, IL-6, and TNF-α) have also been related to OHSS (78,84–86). Proteomic data using human blood and follicular fluid showed that patients with a high risk of OHSS tended to have higher expression levels of proteins related to inflammation, complement and coagulation cascades, and cell adhesion (87,88). Consistent with these previous findings, our RNA-seq data discovered that follicles cultured with excessive FSH had significantly higher expression levels of *Ptgs2*, and a number of proinflammatory genes (Figure 7D), suggesting that there are other potential risks factors of OHSS induced by high doses of FSH, such as the high induction of LHCGR, PTGS2, and related PEG2 production.

The transcriptomic dose-response analysis of FSH enabled us to identify a number of new genes and signal transduction pathways that may critically regulate gonadotropin-dependent folliculogenesis. Compared to follicles cultured with 5 mIU/mL FSH on day 4, *Krt19* was the most up-regulated gene in follicles cultured with 10-30 mIU/mL FSH (Figure 4C). *Krt19* has been used as a granulosa cell marker in stem cell-differentiated granulosa cell-like cells (89,90). Although the role of *Krt19* in folliculogenesis is unknown, its encoded protein has been shown to act as a cytoplasmic intermediate filament to control the structural rigidity and scaffolding in liver hepatocytes and hair matrix epithelial cells (91), suggesting that FSH-induced *Krt19* may play a similar structural role in granulosa cells. Among the top 10 DEGs between day 6 vs day 4, *Cldn3*, *Klk1*, and *Scarf1* were up-regulated in all FSH concentration groups (Figure 4C). All three genes have also been reported to regulate the tight junction and *de novo* angiogenesis (92–94), suggesting their crucial roles in the establishment of tight junction and angiogenesis during FSH-stimulated follicle maturation.

*Nts*, a gene encoding neuropeptide neurotensin, is among the top 10 up-regulated genes in both 20 and 30 mIU/mL FSH groups (Figure 4C). *Nts* has been reported as a paracrine factor to regulate follicle rupture during LH-induced ovulation by stimulating the focal adhesion (95). Indeed, several other ovulatory genes encoding cytokine receptors, such as *Cx3cr1* and *Tnfrsf9*, were also among the top 10 up-regulated genes in the 30 vs 5 mIU/mL FSH comparison groups (Figure 4C). These genes have been implicated in inflammation during ovulation (96). Using the same eIVFG system, our published RNA-seq data also discovered a significant up-regulation of *Nts* and *Tnfrsf9* by 4- and 195-fold, respectively, during *ex vivo* ovulation (45). These results suggest that excessive FSH promotes the expression of proinflammatory genes; despite that the induction of these transcripts is not sufficient to trigger ovulation, they may cause ovarian inflammation.

Both folliculogenesis and oogenesis highly rely on metabolic processes with bidirectional communications between oocyte and its surrounding granulosa cells (97). Compared to the subthreshold level of FSH at 5 mIU/mL, FSH at >=10 mlU/mL significantly altered several metabolic pathways, primarily including glycolysis and related starch and sucrose metabolism and biosynthesis/metabolism of amino acid (glycine, serine, threonine, and arginine, Figure 6). These results are consistent with previous studies which showed the activation of metabolic signals during gonadotropin-dependent follicle maturation in mice (98). Glycolysis, a critical source of Adenosine triphosphate (ATP), occurs in granulosa cells to support cell proliferation and in oocytes to support oocyte development. Transport of granulosa cell glycolysis intermediates, such as pyruvate and lactate, to oocytes is also crucial for generating energy to underpin oogenesis (97,99,100). The oocyte-derived paracrine factors, such as growth differentiation factor 9 (GDF9), bone morphogenetic protein 15 (BMP15), and fibroblast growth factor 8 (FGF8), in turn promote glycolysis in cumulus granulosa cells (101,102). In our study, the transcriptional induction of glycolytic enzyme-related genes, such as *Aldob*, *Aldoc*, and *Eno2*, exhibited a FSH concentration-dependent manner (Figure 6C) (33). A previous study reported the over-accumulation of glycolytic intermediates induced by excessive FSH, such as lactate, which may cause pyknotic nuclei surrounding oocytes, an early sign of follicle atresia (50). Interestingly, our single-oocyte RNA-seq data showed that 30 mIU/mL FSH reduced the expression of *Fgf8* compared to 10 mIU/mL (Suppl. Table 8), suggesting a potential disruption of glycolysis during oogenesis. Our results also revealed that excessive 30 mIU/mL FSH significantly promoted the lipid metabolism in follicles compared to 10 mIU/mL FSH (Figure 7). Lipid metabolism is crucial for steroidogenesis in granulosa cells as well as the energy production for oocyte maturation and luteinization (98,103,104). Treatment with high levels of gonadotropins in a mouse superovulation model caused more lipid accumulation in 2-cell staged embryos, potentially attributed to the increased P4 due to enhanced cholesterol production (105). Cortisol has been identified as an upstream regulator of lipid metabolism in follicular cells (106). Our results showed that *Hsd11b1*, a gene encoding hydroxysteroid 11-beta dehydrogenase 1 (HSD11B1) for cortisol synthesis, increased by 3-fold in follicles treated with 30 mIU/mL vs 10 mIU/mL FSH (Suppl. Table 3). However, the potential impact of excessive FSH-induced lipid metabolism changes on oocyte quality requires further investigations.

Previous studies investigating the effects of high doses of FSH during ovarian stimulation on oocyte quality showed inconsistent results. Several studies reported detrimental effects of excessive FSH on oocyte quality, but others found no associations (107–110). For instance, the administration of high doses of FSH in IVF patients has been linked to oocyte brown zona pellucida thickness, embryo aneuploidy, and reduced pregnancy rates (107,108). In contrast, other studies reported no adverse impacts on oocyte quality and related embryo development and pregnancy outcomes, as evidenced by the comparable rates of fertilization and pregnancy success between different FSH dosing groups (109,110). Similar to our single-oocyte RNA-seq results, oocytes from a mouse superovulation model have been shown to have similar transcriptomic profiles compared to oocytes from naturally cycling mice (111). The contradictory findings might be caused by variations in IVF patients, ovarian stimulation protocols, species differences, and evaluation methods. So far, most DEGs identified in our single-oocyte RAN-seq analysis have not been implicated in oogenesis (Figure 10). *Frk*, a gene encoding a protein belonging to the TYR family of protein kinases, increased for 2.1-fold by 30 mIU/mL FSH. Despite *Frk* encoding protein, Fyn related Src family tyrosine kinase, has not been established to regulate oogenesis, other Src family tyrosine kinases, such as SRC, FYN, and YES, may contribute to oocyte maturation and fertilization.

Maternal effect genes accumulate during oogenesis, which are critical for the acquisition of oocyte developmental competence, epigenetic reprogramming, imprinting, maternal to embryonic control transition, and early embryogenesis (112). Our results showed that all 70 maternal effect genes identified to date had comparable expression levels in oocytes from follicles treated with 10 and 30 mIU/mL FSH (Figure 10), indicating that a high dose of FSH may not affect oocyte maternal effect genes. The administration of high dose of FSH in a mouse superovulation model has been shown to affect several maternal imprinted genes (113) established during oogenesis (114,115), suggesting that it is worth investigating the effects of excessive FSH on epigenetic modifications in oocytes. The mitochondria in oocytes primarily regulate energy metabolism and Ca^2+^ homeostasis to sustain chromosome stability (116,117). Dysfunctional mitochondria, such as reduced mitochondrial ATP, has been shown to cause spindle disassembly in mouse oocytes (118). Our results showed that the expression of several genes contributing to mitochondrial structure and functions were downregulated by 30 mIU/mL FSH, including those encoding the inner mitochondrial membrane protein (*Slc25a1*, *Tomm40, Gpx4*) and NAD activity/energy generation related protein (*Aldh2*, *Aldh7a1*, *Idh3b*). More interestingly, all these genes have been implicated in oocyte viability and maturation (119–122), suggesting that the reduction of mitochondrial genes caused by excessive levels of FSH may compromise the quality of oocytes and subsequent reproductive outcomes. Since both the comparable oocyte maternal effect genes and reduced mitochondrion-related genes discovered in our study are at the transcriptional level, further studies comparing the oocyte proteomics and developmental outcomes are needed (111).

In conclusion, using a functional and transcriptomic FSH dose-response analysis of an *ex vivo* mouse folliculogenesis and oogenesis system, we demonstrate that there is a minimum FSH threshold required to support gonadotropin-dependent terminal follicle maturation and related ovarian and reproductive functions. Moreover, there exists an FSH upper-bound threshold which is sufficient to stimulate the maturation of all follicles without inducing ovarian defects. Excessive FSH above this upper-bound threshold may heighten the risks of premature luteinization, hyperandrogenism, ovarian inflammation, and reduce oocyte energy metabolism, which potentially compromises follicle and oocyte quality and result in adverse female reproductive outcomes. These novel findings improved our understanding of gonadotropin-dependent follicle maturation at the molecular level and provide crucial insights into optimizing controlled ovarian stimulation protocols in ART to achieve better and safer IVF outcomes.

## Supporting information

Supplemental Materials, and will be used for the link to the file on the preprint site.

Supplemental Table, and will be used for the link to the file on the preprint site.

## Author Contributions

T. Zhan contributed to the experimental design, data collection and analysis, and manuscript writing; J. Zhang, and Y. Zhang contributed to the data collection and analysis; Q. Zhao and NC. Douglas contributed to the manuscript writing; and S. Xiao and Q. Zhang conceived the project, designed experiments, interpreted data, wrote the manuscript, and provided final approval of the manuscript.

## Data Availability

Associated RNA-seq data have been deposited at the Gene Expression Omnibus (GEO:GSE249196) (https://www.ncbi.nlm.nih.gov/geo/query/acc.cgi?acc=GSE249196).

