## Supplemental Materials, and will be used for the link to the file on the preprint site. for "Dose-response functional and transcriptomic effects of follicle-stimulating hormone on *ex vivo* mouse folliculogenesis"

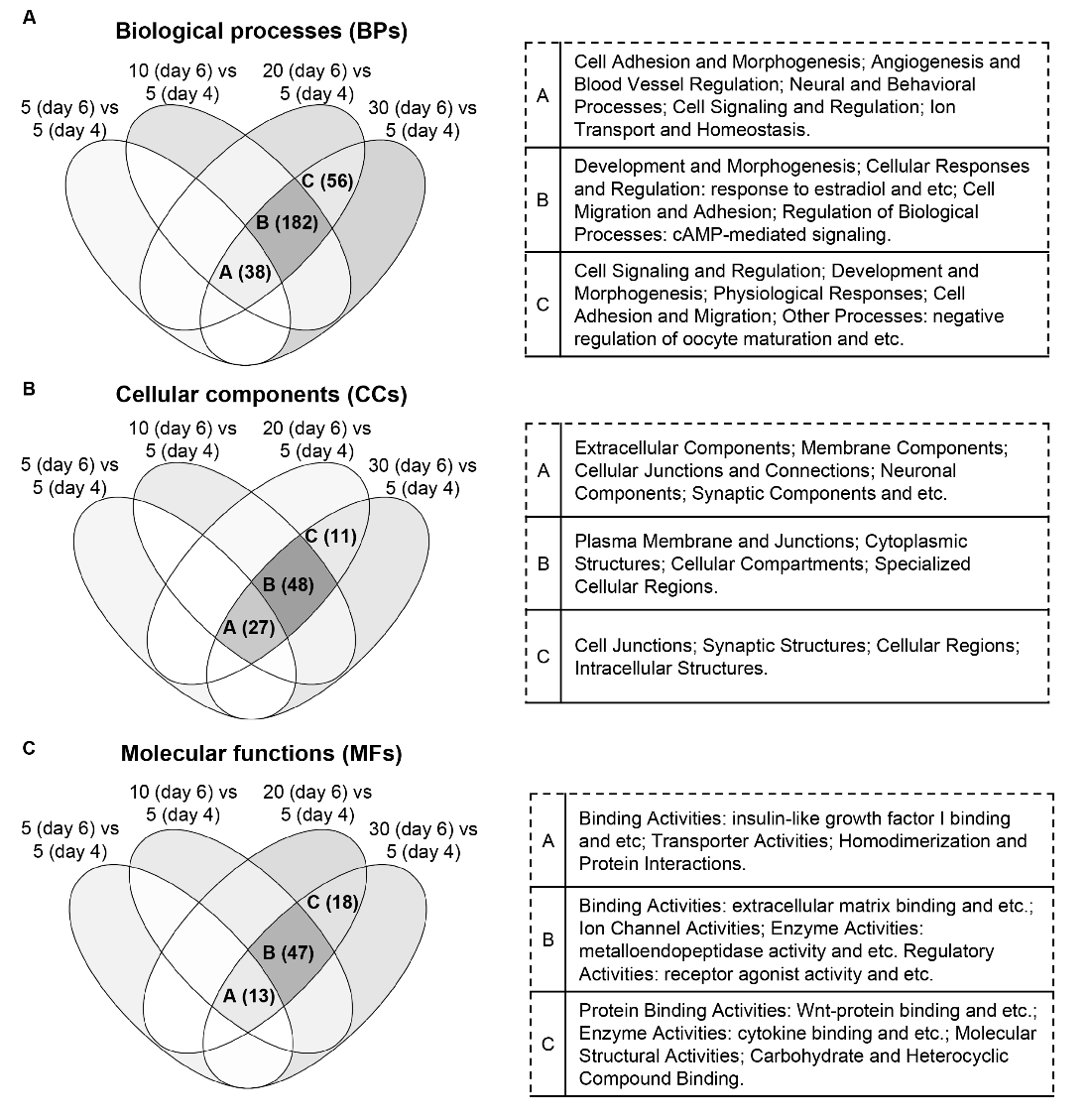

**Supplementary Figure 1.** Venn diagram of biological processes (BPs), cellular components (CCs) and molecular functions (MFs) enriched by DEGs in follicles treated with FSH at 5, 10, 20 and 30 mIU/mL on day 6 compared to follicles on day 4.

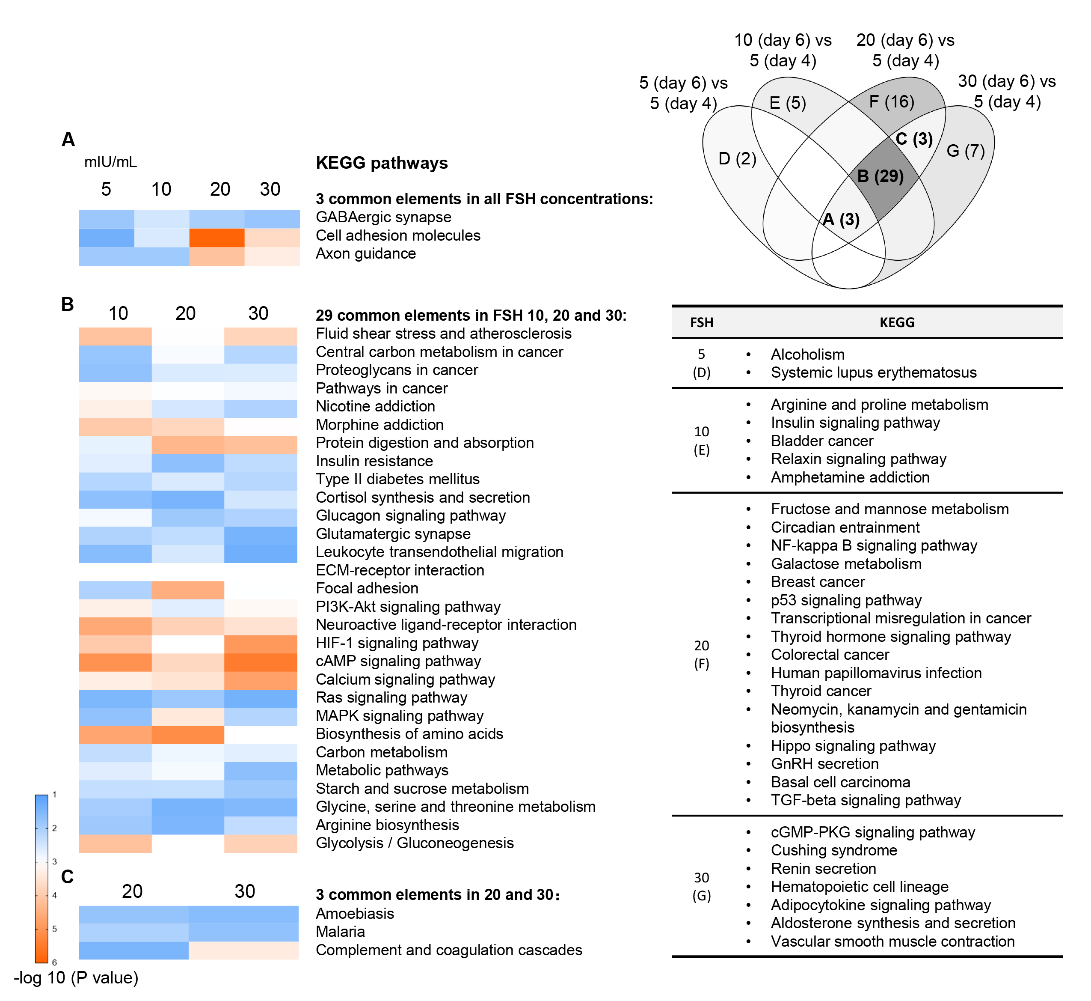

**Supplementary Figure 2.** Comparison of KEGG pathways in follicles treated with FSH at 5, 10, 20 and 30 mIU/mL on day 6 compared to follicles on day 4. (A) Common KEGG pathways enriched in all FSH concentration groups. (B) Common KEGG pathways only enriched in FSH of 10, 20 and 30 mIU/mL. (C) Common KEGG pathways only enriched in FSH of 20 and 30 mIU/mL.

**Supplemental Table 1.** Primer sequences of examined genes by RT-qPCR

| **Gene** | **Forward Primer (5’-3’)** | **Reverse Primer (5’-3’)** |
| --- | --- | --- |
| *Gapdh* | CATCACTGCCACCCAGAAGACTG | ATGCCAGTGAGCTTCCCGTTCAG |
| *Star* | AAGCTGTGTGCTGGAAGCTC | CTTCCAGTTGAGAACCAAGCAG |
| *Cyp11a1* | TCAAAGCCAGCATCAAGGAGA | TGGCAAAGCTAGCCACCTGTA |
| *Cyp17a1* | CCTGATACGAGGCACTTCTCG | CCAGGACCCAAGTGTGTTCT |
| *Hsd3b1* | AGTGATGGAAAAAGGGCAGGT | GCAAGTTTGTGAGTGGGTTAG |
| *Hsd17b1* | ACTGTGCCAGCAAGTTTGCG | AAGCGGTTCGTGGAGAAGTAG |
| *Cyp19a1* | CATGGTCCCGGAAACTGTGA | GTAGTAGTTGCAGGCACTTC |
| *Fshr* | GAGGCAGATGTGTTCTCCAACC | TCGGAGACTGGGAAGATTCTGG |
| *Lhcgr* | GACGCTAATCTCGCTGGAGT | GGCCTGCAATTTGGTGGAAG |
| *Pappa* | CAGAAAGCCAGCACCTGTAGCT | GGCAAAGGTCACATGCTGATCC |
| *Inhba* | GGAGATAGAGGACGACATTGGC | ACGCTCCACTACTGACAGGTCA |
| *Nppc* | GGTCTGGGATGTTAGTGCAGCTA | TAAAAGCCACATTGCGTTGGA |
| *Amh* | CAGGCCCTGTTAGTGCTATACCCT | GAAGTCCACGGTTAGCACCAAA |
| *Vegfa* | AGGGTCAAAAACGAAAGCGC | CACATCTGCAAGTACGTTCG |
| *Vegfc* | GAGGTCAAGGCTTTTGAAGGC | CTGTCCTGGTATTGAGGGTGG |
| *Areg* | CACAGCGAGGATGACAAGGA | ATCGTTTCCAAAGGTGCACTG |
| *Ereg* | GCATCCCAGGAGAATCCGAG | GTGTAGCCCACTTCACATCTGC |
| *Btc* | AAACCCACTTCTCTCGGTGC | GCCTTTCTCACAGATGCAGG |
| *Has2* | GCCGGTCGTCTCAAATTCATC | ACCTCTCACAATGCATCTTGTTC |
| *Tnfaip6* | GCTACAACCCACATGCAAAGG | CTGACCGTACTTGAGCCGAA |
| *Ptgs2* | TTCTTTGCCCAGCACTTCAC | CACCTCTCCACCAATGACCTGA |
| *Pgr* | TTAAGAGGGCAATGGAAGGGCA | TTTCTCAGACGACATGCTGGG |
| *Runx1* | CCTTCAGGAGAGGTGCGTTT | CTCGTGCTGGCATCTCTCAT |
| *Runx2* | CCTGAACTCTGCACCAAGTCCT | TCATCTGGCTCAGATAGGAGGG |
| *Il6* | CCGGAGAGGAGACTTCACAG | TCCACGATTTCCCAGAGA |
| *Ccl2* | CTTCTGGGCCTGCTGTTCA | CCAGCCTACTCATTGGGATCA |
| *Cxcl1* | TCCAGAGCTTGAAGGTGTTGCC | AACCAAGGGAGCTTCAGGGTCA |
| *Adamts1* | TTGAATGGTGTGAGTGGCGA | CCATCAAACATTCCCCGTGTC |
| *Plat* | AAGAAGCAAGCACTCTCGGG | TCATCTCTGCAGGTCGCTCT |
| *Plau* | CATCCAGTCCTTGCGTGTCT | CCAAGTACACTGCCACCTTCA |

**Supplemental Table 2.** Ovarian steroidogenesis and follicle maturation-related genes and their functions

| **Gene** | **Description** | **Functions and references** |
| --- | --- | --- |
| *Star* | Steroidogenic acute regulatory protein | Expressed in theca cells. Transfer of cholesterol from the outer to the inner mitochondrial membrane (1). |
| *Cyp11a1* | Cytochrome P450 family 11 subfamily A member 1 | Conversion of cholesterol to pregnenolone (2) |
| *Cyp17a1* | Cytochrome P450 family 17 subfamily A member 1 | Expressed in theca cells. Hydroxylation progesterone/pregnenolone at the 17 carbon, and conversion of 21-carbon steroids to androstenedione or dehydroepiandrosterone (DHEA) (3) |
| *Hsd3b1* | Hydroxy-delta-5-steroid dehydrogenase, 3 beta- and steroid delta-isomerase 1 | Conversion of pregnenolone to progesterone, 17alpha-hydroxypregnenolone to 17alpha-hydroxyprogesterone, (DHEA) to 4-androstenedione (4) |
| *Hsd17b1* | Hydroxysteroid 17-beta dehydrogenase 1 | Conversion of androstenedione to testosterone, estrone to estradiol (5) |
| *Cyp19a1* | Cytochrome P450 family 19 subfamily A member 1 | Expressed in GCs. Encoding aromatase responsible for the aromatization of androgens into estradiol (6) |
| *Fshr* | Follicle stimulating hormone receptor | Transmembrane receptor of FSH for activating FSH target signaling and genes (7,8) |
| *Lhcgr* | Luteinizing hormone/chorionic gonadotropin receptor | Transmembrane receptor of LH for regulation of follicular maturation and ovulation (9,10) |
| *Pappa* | Pregnancy-associated plasma protein A | IGFBP protease for activation of IGF pathway and regulation of folliculogenesis (11) |
| *Inhba* | Inhibin subunit beta A | Synthesis and secretion of ovarian peptide hormones of inhibin and activin  and regulation of folliculogenesis (12-15) |
| *Nppc* | Natriuretic Peptide C | Granulosa cell ligand for producing cGMP and maintaining oocyte meiotic arrest (16) |
| *Amh* | Anti-müllerian hormone | An marker for ovarian reserve with the inhibitory effect on folliculogenesis (17) |

**Supplemental Table 7.** Ovulation-related genes and their functions

| **Gene** | **Description** | **Ovulatory functions and references** |
| --- | --- | --- |
| *Vegfa* | Vascular endothelial growth factor-A | Activation of vascularization for follicle ovulation (18) |
| *Vegfc* | Vascular endothelial growth factor-C |  |
| *Areg* | Amphiregulin | Activation of EGF signaling to regulate cumulus expansion, oocyte maturation, and follicle rupture (19-21) |
| *Ereg* | Epiregulin |  |
| *Btc* | Betacellulin |  |
| *Has2* | Hyaluronan synthase 2 | Cumulus cell expansion regulator (22) |
| *Tnfaip6* | TNF alpha induced protein 6 | Cumulus cell expansion regulator (23,24) |
| *Ptgs2* | Prostaglandin-endoperoxide synthase 2 | Cumulus cell expansion regulator (25) |
| *Pgr* | Progesterone receptor | Transcription factor (26,27) |
| *Runx1* | RUNX Family Transcription Factor 1 | Transcription factor (28) |
| *Runx2* | RUNX Family Transcription Factor 2 | Transcription factor (29) |
| *Il6* | Interleukin 6 | Proinflammatory factor (30) |
| *Ccl2* | C-C motif chemokine ligand 2 | Proinflammatory factor (31-33) |
| *Cxcl1* | C-X-C Motif Chemokine Ligand 1 | Proinflammatory factor (34) |
| *Adamts1* | A disintegrin and metalloproteinase with thrombospondin motifs 1 | Proteolysis, ECM remodeling, and follicle rupture (26,35) |
| *Plau* | Plasminogen activator, urokinase | Proteolysis, ECM remodeling, and follicle rupture (36,37) |
| *Plat* | Plasminogen activator, tissue type | Proteolysis, ECM remodeling, and follicle rupture(26,35-37) |
